# Neuropeptide Dynamics Coordinate Layered Plasticity Mechanisms Adapting Drosophila Circadian Behavior to Changing Environment

**DOI:** 10.1101/2024.10.15.618497

**Authors:** Abhishek Chatterjee, Joydeep De, Béatrice Martin, Elisabeth Chélot, François Rouyer

## Abstract

The Drosophila brain contains distinct sets of circadian oscillators responsible for generating the morning and evening bouts of locomotor activity, giving rise to a bimodal rest-activity pattern in light-dark cycles. We lack a mechanistic understanding of how environmental changes reshape this daily profile of rest-activity pattern. Here, we uncover a seasonal switch mechanism that remodels the evening bout of activity. Under summer-like conditions, an environment favored by fruit flies in temperate climates, levels of the PDF neuropeptide diminish, triggering a cascade. Lowered PDFR signaling disinhibits GSK3/SGG to advance the evening output. Upon sensing PDF loss, the neural activity weakens in the DN1p-SIF*a* circuit, responsible for promoting afternoon rest; leading to an earlier appearance of the evening peak. At the same time, the functional connections from DN1p to LNd oscillators strengthen, consequently handing over the evening pacemaker role to the DN1ps. Taken together, our findings elucidate how environment-induced changes in PDFR signaling tip the balanced output of the clock network, aligning daily rhythms with seasonal time. Neuropeptide-driven parallel adjustment of clock circuitry and clock protein functioning likely represents a conserved strategy across animal species, enabling them to adapt their daily behavior to seasonal changes throughout the year.

## Introduction

Nearly every organism possesses a circadian clock that organizes physiological and behavioral processes according to the time of the day. The working mechanism of eukaryotic clocks is centered on transcriptional feedback loops, with post-transcriptional controls setting the clocks’ period close to 24 hours ^1^. While the period of the external day-night cycles is characteristically invariant, the day-to-night ratio varies seasonally in most places on Earth. The clock-driven rhythms have therefore evolved to manifest a stable period, but a labile waveform and phase. In mammals, encoding of daylength relies on the plasticity of the circadian waveform in the mammalian suprachiasmatic nucleus (SCN), but the network changes underlying waveform plasticity remain enigmatic ^2^. The 150 clock neurons of the *Drosophila* brain are particularly suited for network analysis, being organized into functionally and anatomically segregated, variably coupled oscillator nodes ^3,4^.

In the fruit fly, the conserved biochemical program of the circadian oscillator, at its core, involves CLOCK (CLK) – CYCLE (CYC) –driven transcription of *period (per)* and *timeless (tim)* genes ^1^. The PERIOD (PER) and TIMELESS (TIM) proteins associate to form complexes that repress CLK-CYC activity. A series of post-transcriptional modifications control the subcellular location, half-life and transcriptional potency of these protein complexes. Light and temperature inflict state-change of the dynamic clock program, mostly by affecting the stability of PER-TIM ^5–7^. Consequently, light and temperature changes synchronize the brain clock with the external day-night cycles (entrainment) ^8,9^. The external time cues (zeitgeber) also tune the crosstalk between oscillator nodes, and selectively gate the oscillator’s overt output ^10,11^. The multiscale effect of light and temperature on the clock guarantees exquisite malleability of the rhythms’ phase and waveform across phylogeny.

The daily rest-activity pattern of *Drosophila* is bimodal ^12^ as it is in humans ^13,14^. Dawn-active morning oscillator (MO) and dusk-active evening oscillator (EO) neurons in the *Drosophila* brain produce the two daily peaks of locomotor activity in anticipation of day/night transition ^15–18^. There is support for the general idea that the circadian clock is instrumental in seasonal time measurement ^19,20^. In the rodent suprachiasmatic nucleus (SCN), the phase-angles of individual oscillator neurons diverge as the day gets longer, wherein the VIP neuropeptide (Vasoactive Intestinal Peptide) plays a key role in adapting the behavioral activity profile to photoperiodic changes ^21–24^.

A similar mechanism implicating inter-oscillator phase-gap was invoked to explain fly’s behavioral response to daylength change ^17^. Seasonal variation in the daily pattern of light and temperature changes necessitates flexibility in the timing of the two activity peaks –two peaks moving closer or apart depending on the prevailing daylength and temperature, and the morning and evening oscillators do show different responses to light and temperature ^17,25–27^. *Drosophila*’s activity peaks and their source oscillators, however, have a markedly limited ability to track the dawn and dusk transitions across different photoperiods ^28,29^, which challenges the long-standing idea that seasonal time is encoded by photoperiod-dependent shifts in inter-oscillator phase-angle^30^.

In addition to the phase changes, observations in semi-natural conditions revealed a dramatic seasonal change in waveform: hot, dry, and intensely bright summer days induce an afternoon bout of activity in tropical and Mediterranean zones ^31–33^. Conversely, waveform plasticity was also observed in subarctic Drosophilids that experience unusually long yet mild summer days ^34–36^. Under these polar conditions, the evening peak becomes more pronounced, whereas in the standard laboratory setting characterized by high-amplitude 12h:12h light-dark (LD) cycles, similar to equinox days–the evening peak is only as strong as the morning one, and remains tied to the dusk transition. Notably absent is our understanding of the plasticity mechanisms that enable the monitoring of seasonal changes and their translation into behavioral adjustments, especially under the integrative influence of variations in daylength, light, and temperature across seasons.

The neuropeptide Pigment-Dispersing Factor (PDF), synthesized by morning cells (MO), is involved in developmental and physiological responses to photoperiod ^37–40^. Downstream of the clock neuronal network, PDFR signaling promotes degradation of the EYES ABSENT (EYA) phosphatase in insulin-producing cells causing female reproductive dormancy on winter days ^38,41^. PDF is also important for the photoperiodic adjustment of the fly’s evening activity peak ^42–48^. Nonetheless, the tunable organizational changes that enable clock outputs including PDF to drive seasonal adaptation of the daily rest-activity pattern remain elusive.

In this study, we demonstrate that PDF-mediated communication among clock neurons in the Drosophila brain edits the network organogram. By reducing the neural activity of a locomotor-suppressive output circuit, a larger evening peak is produced on summer days. This tunable balance between locomotor-suppressive and –promotive oscillators, set by neuropeptide dynamics, could allow the multi-oscillator mammalian clock to adaptively mold behavioral waveform and phase.

## Results

### Summer-like conditions induce a drop of PDF levels in cosmopolitan Drosophila species

Wild-type *D. melanogaster* flies start to build evening anticipatory activity 3-4 hours before lights-OFF in equinox days of standard laboratory conditions (LD 12:12, 25°C). To test flies in conditions that better correspond to their natural environment in temperate climates, we used long days and increased temperature. Owing to the formation of new leaves, light penetration and availability are strongly decreased on summer days in woodlands and fruit orchards that constitute the natural habitat of Drosophila (see Mat. and Meth.). Moreover, flies actively seek low light intensities, particularly at warmer temperatures ^49–51^. We thus applied lower light intensity in our laboratory-simulated summer days (LD 16:8 low light, 28°C).

A broader peak starting much earlier before lights-OFF was observed in temperate summer-like conditions (Fig. 1A, Fig. S1A). Since the absence of PDF advances the evening activity in laboratory LD 12:12 cycles ^42^, we asked whether environmental cues could be encoded by PDF signaling itself. In the absence of PDF signaling, a surprisingly small effect of the environmental conditions was observed on evening activity, barring a modest increase in phase advancement (Fig. 1A, Fig. S1A). This suggested that the repatterning of daily activity in summer-like conditions could be brought about by a change of PDF signaling. We thus quantified PDF levels under the two different conditions. In summer-like conditions, PDF peptide and *Pdf* mRNA were greatly diminished in the two subsets of ventral lateral neurons (LNvs) (Fig. 1B). In parallel, expression of the PDF receptor gene *(Pdfr)* also plummeted in the LNds and DN1ps (Fig. 1C), which play key role in the control of evening activity ^10,15,16,52^. Interestingly, the absence of PDF decreased *Pdfr* expression and the absence of PDFR decreased PDF levels (Fig. S1B). Therefore, we inferred that temperate summer-like conditions are marked by a coordinated drop in levels of PDF and its receptor.

**Figure 1.**
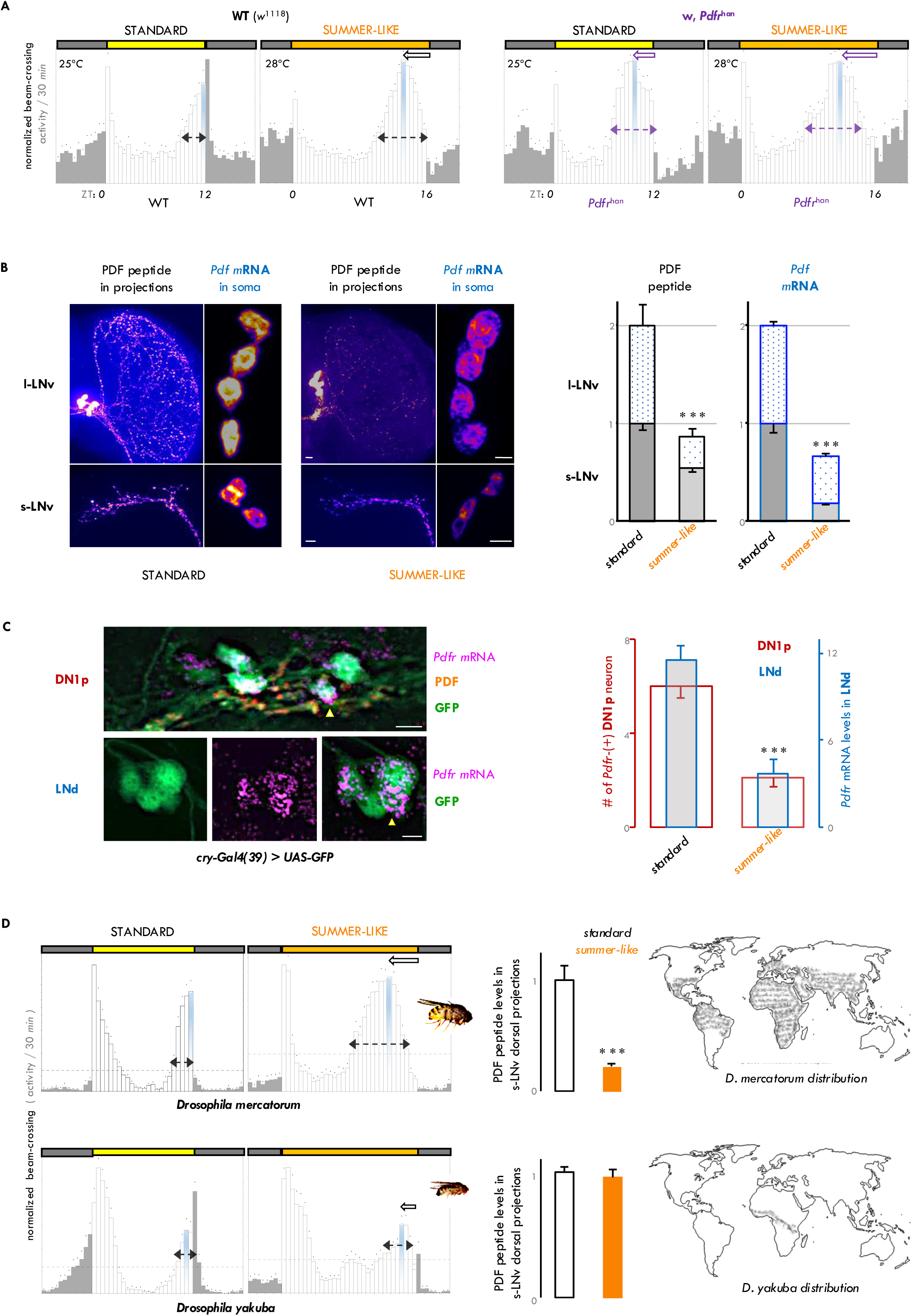
Broad and advanced evening peak of summer days is linked to seasonally weakened PDF signaling tone in cosmopolitan drosophilids. **A**. Normalized average daily locomotor activity profiles (eductions) of wild-type (WT: *w*^1118^) and *Pdfr*^han^ mutant *Drosophila melanogaster* flies in equinox and summer conditions. The white bars represent the activity during the light phase (colored box) and the grey bars represent the activity in the dark period (grey box), the dots represent the s.e.m. The peak of evening (E) activity is marked in blue. Dotted arrows denote the width of the E-peak and solid arrows denote advancement of the E-peak’s phase. Photoperiod and temperature conditions are mentioned on the eductions. n=10, 16, 16, 19 flies from left to right. **B**. PDF peptide and mRNA are quantified in s-LNv and l-LNv neurons at the midpoint of the photoperiod in equinox and summer conditions. Both PDF peptide in projections and mRNA in the soma were diminished in summer conditions (n ≥ 14 brain hemispheres; p < 0.0001 after unpaired two-tailed Student’s *t*-test; error bar = s.e.m.). **C**. *Pdfr* mRNA detected by RNAscope in a subset of the LNd and DN1p neurons. On summer days, *Pdfr* level was lower in the LNds, and fewer DN1ps could be co-labeled with the *Pdfr* probe (n ≥ 10 brain hemispheres; p < 0.0001 after unpaired two-tailed Student’s *t*-test; error bar = s.e.m.). **D**. Cosmopolitan *D. mercatorum* is distributed widely as opposed to Afro-tropical *D. yakuba.* Normalized average daily profile of *D. mercatorum* and *D. yakuba* locomotor activity in equinox and summer conditions (n=15-17 flies). Like *D. melanogaster* and unlike *D. yakuba*, cosmopolitan *D. mercatorum* shows broadened and advanced evening (E) activity in summer, correlated with the lowering of PDF peptide level in the s-LNv dorsal termini (n ≥ 8 brain hemispheres; p < 0.0001 after unpaired two-tailed Student’s *t*-test; error bar = s.e.m.).

On summer days, a broad afternoon peak is also displayed by many high-latitude-restricted Drosophila species, even if the photoperiodic adaptation strategy varies among these northern flies ^7,34–36,53,54^. In light of the changes in the activity pattern and PDF levels that we observed in *D. melanogaster*, we asked how other drosophilids adapt to summer-like conditions. Distantly related *D. mercatorum*, cosmopolitan in distribution like *D. melanogaster*, exhibited pronounced broadening of the evening activity and phase-advancement of the evening peak in summer-like conditions (Fig. 1D, Fig. S1C) ^54^. In contrast, the equatorial *D. yakuba* had restricted and delayed evening activity when confronted with high-latitude summer-like conditions (Fig. 1D, Fig. S1C) ^55^. As observed in *D. melanogaster* on temperate summer days, PDF levels in the s-LNv axonal termini strongly decreased in *D. mercantorum*; conversely, PDF levels did not change between standard and summer-like conditions in the Afrotropical *D. yakuba* flies (Fig. 1D). *D. malerkotliana* flies which has an intermediate latitudinal range, also underwent PDF decrease in summer days, accompanied by the pronouncement of its evening peak (Fig. S1C-D) ^56^. In temperate summer-like conditions, a broad early evening peak of activity thus appears to be correlated with a decrease of PDF signaling in cosmopolitan Drosophila species.

### A PDF-PKA pathway delays evening activity by inhibiting SGG-dependent TIM degradation

To understand how weakened PDFR signaling on summer days broadens and advances the evening peak, we sought to determine whether the loss of PDF speeds up the evening oscillator in the CRY-positive PDF-negative LNs (LN-EO). In *Pdf*^0^ flies, TIM and PER proteins normally accumulated until the middle of the night, but then, TIM levels and subsequently PER levels decreased significantly faster than in the wild-type (Fig. 2A). This suggested that enhanced PER/TIM degradation at the end of the night advances the evening peak in the absence of PDF.

**Figure 2.**
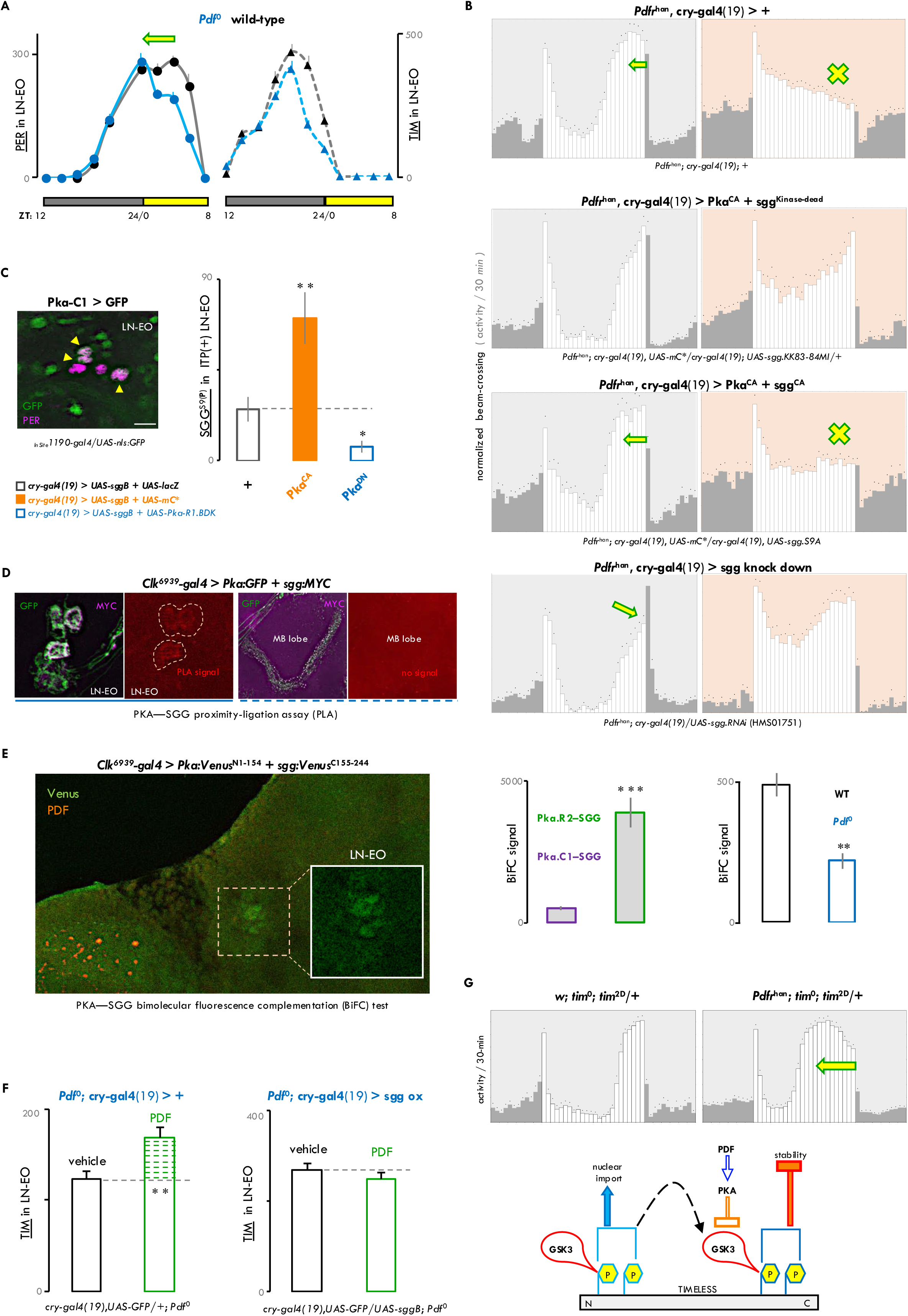
PDF-regulated PKA signaling working through SGG/GSK3 impinges on TIM to delay the LN-EO clockwork. **A**. Immunostaining against PERIOD (PER) and TIMELESS (TIM) proteins in *w;;Pdf*^0^ and wild-type (*w*^1118^) flies in 12:12 light-dark (LD) cycle. The entire dark phase and the first 8 hours of the light phase are shown. TIM, and subsequently, PER levels diminished significantly faster (p<0.001 by two-way ANOVA) towards the end of the night in *Pdf*^0^ flies. 8-14 brain hemispheres were analyzed for each time point. error bar = s.e.m. **B**. Normalized average daily profiles of locomotor activity in LD 12:12 (left) and red-light LD, *i.e.*, RD 12:12 (right) of *Pdfr*^han^ flies with different combinations of PKA and/or SGG modifications in the LNs. In *Pdfr^han^*flies, constitutively active PKA (Pka^CA^) or *sgg* knockdown rescued the advanced E-peak of LD as well as the E-activity loss of RD. Co-expression of the constitutively active forms of both PKA and SGG (sgg^CA^) cancelled this rescue, reverting to the *Pdfr^han^* phenotype of LD and RD. **C**. (Left) A subset of the LNds show prominent expression of the C1-catalytic subunit gene of the PKA holoenzyme. (Right) In the soma of the ITP-expressing LN-EO neurons, levels of S9-phosphorylated, hence catalytically inhibited SGG, increased (p < 0.01 by *t*-test) upon PKA over-activation (Pka^CA^), while PKA downregulation by dominant-negative regulatory subunit (R1.BDK) expression reduced S9-phosphorylated SGG level (p < 0.05 by *t*-test). Brains were dissected around ZT6. (n ≥ 14 brain hemispheres; error bar = s.e.m.). **D**. Positive signal (red) in the *in-situ* Proximity Ligation Assay (PLA) suggests close (≤ 40 nm) juxtaposition between MYC-tagged SGG molecules (purple) and GFP-tagged PKA holoenzyme (green) in the soma of the LN-EO neurons but not in the mushroom body (MB lobes) where both the molecules are still present. Samples were analyzed at ZT2-4. **E**. (Left) Reconstitution of native *Venus* signal (green) through Bimolecular Fluorescence Complementation (BiFC) between PKA-C1 and SGG molecules in the LN-EO. Successful BiFC is contingent upon close proximity (≤ 7 nm) between the two molecules, consistent with the idea that they either directly interact or are part of the same macromolecular protein complex. (Right) Stronger BiFC signal with SGG was elicited by the R2 regulatory subunit of PKA compared to its C1 catalytic subunit (p < 0.001 by *t*-test). Interaction between PKA-C1 and SGG was weaker (p < 0.01 by *t*-test) in the absence of PDF, *i.e.*, the upstream activator of PKA signaling in the LN-EO. (n ≥ 10 brain hemispheres; error bar = s.e.m.). **F**. At ZT14, exposure of *ex-vivo* brains to 0.1 mM PDF for 3.5 hr stabilized TIM (p < 0.01 by *t*-test), in the LN-EO neurons of *Pdf*^01^ flies. No significant (p = 0.17 by *t*-test) effect of PDF on TIM stability was apparent in the same cells when SGG was overexpressed in *Pdf*^01^ background. (n ≥ 10 brain hemispheres; error bar = s.e.m.). **G**. (Top) A variant of TIM (TIM^2D^) bearing phospho-mimetic substitutions of 2 amino acids otherwise phosphorylated by SGG, triggered strong interaction with *Pdfr^han^* mutation manifested in dramatic advancement of the E-peak phase in LD. (Bottom) Our working model suggests that PDF-activated PKA signaling inhibits SGG/GSK3 kinase activity, thereby slowing down TIM turnover. Low PDF on summer days should then relieve the inhibition on SGG kinase and hence hasten TIM turnover.

Protein Kinase A (PKA) signaling is involved in the PDF-dependent phasing of the evening activity ^57^. In PDF-negative neurons, PKA promotes PER and TIM stability ^57–59^ but the mechanism is not known. The SHAGGY (SGG) kinase accelerates the clock by phosphorylating TIM ^60,61^ and SGG’s mammalian ortholog GSK3 is inhibited through Ser-9 phosphorylation by PKA ^62,63^. We thus hypothesized that SGG/GSK3 could be inhibited by PKA in the PDFR signaling pathway. Constitutively active PKA (PKA^CA^) rescues the advanced evening peak of *Pdfr*^han^ mutants ^57^ and we predicted that simultaneously increasing SGG activity would inhibit this rescue. Indeed, constitutively active SGG^CA^ nullified the PKA^CA^-dependent rescue, whereas SGG knockdown rescued the advanced evening peak of *Pdfr*^han^ flies having low PKA activity (Fig. 2B, left). The PDFR-PKA signaling pathway thus appears to delay evening activity by counteracting SGG. Under red-light LD (RD) cycles that prevent CRY activation, *Pdfr*^han^ mutants lose the evening peak ^44^. We took advantage of this robust, qualitative RD phenotype to demonstrate the link between PDFR, PKA, and SGG. *Pdfr*^han^ mutants expressing constitutively active PKA^CA^ got back the evening peak in RD, but not if SGG^CA^ was simultaneously present (Fig. 2B, right). Furthermore, SGG knockdown restored the evening peak of *Pdfr*^han^ flies in RD (Fig. 2B, right). Altogether, the LD and RD results support that the PDFR-PKA pathway inhibits SGG to properly phase evening activity.

We directly assessed SGG S-9 phosphorylation upon alteration of PKA activity in the LN-EO, which endogenously expresses the *Pka-C1* gene (Fig. 2C, left). PKA^CA^ produced substantially higher levels of S9-phosphorylated-SGG, whereas a dominant negative PKA^DN^ reduced S9 phosphorylation (Fig. 2C, right). We then went on to examine whether PKA phosphorylates SGG by directly engaging with it. *In-situ* proximity ligation assay with GFP-tagged PKA and MYC-tagged SGG revealed evidence for PKA–SGG physical interaction in the cell-body of the LN-EO (Fig. 2D). Bimolecular fluorescence complementation validated cytosolic interaction between PKA and SGG, further demonstrating that SGG interacted more strongly with the R2 regulatory subunit of PKA than with the C1 catalytic subunit (Fig. 2E). PKA-SGG interaction was weaker in *Pdf*^0^ mutant (Fig. 2E). This suggested that PDF signaling facilitates PKA-SGG interaction in the LN-EO, promoting the observed SGG S9-phosphorylation that signifies reduced SGG kinase activity.

Decreased PKA activity destabilizes TIM even in the absence of PER ^57^ and *Pdf*^0^ flies had faster decay of TIM and PER levels, with effect on TIM preceding that of PER (Fig. 2A). This suggested that the PDFR-PKA signaling would inhibit SGG to stabilize TIM. Bath-applied PDF peptide on *ex-vivo Pdf*^0^ brains indeed increased TIM levels at night, at a time when *Pdfr* expression crests in the LN-EO ^64^, and the effect on TIM stabilization was annulled by SGG overexpression (Fig. 2F). To investigate how the PDF-PKA pathway controls SGG-dependent TIM phosphorylation, we used the TIM^2D^ variant that bears 6ncrease-mimetic modifications of two N-terminal amino acids targeted by SGG ^61^. *Tim*^2D^ flies only had a slight advance of the evening peak in LD conditions but displayed a dramatic phase advance in *Pdfr*^han^ background (Fig. 2G, Fig. S2A). This strong, synergistic genetic interaction suggests that PDFR signaling may inhibit SGG phosphorylation at a secondary site whose 6ncrease-occupancy is facilitated by earlier action of SGG on the N-terminal part of TIM (Fig. 2G). Altogether, our results strongly support a model where PDF delays the evening activity through a PKA-SGG kinase-relay that stabilizes TIM at night. Thus, during temperate summer-like conditions, low PDF would induce TIM destabilization, thereby advancing the clock in PDFR-expressing evening cells.

### A new DN1p-resident oscillator is required to broaden the evening activity peak on summer-like conditions

Next, we set out to identify the clock neurons that are involved in the broadening of the evening peak in summer-like conditions. Evening activity is contributed by both the LN-EO and the DN1ps, while the contribution of the DN1ps is distinctly environment-sensitive and gated by PDF signaling ^10,52,65^. We thus hypothesized that a DN1p oscillator could sculpt the daily activity pattern on summer days in response to low PDF, in association with the PDFR-expressing LN-EO. DN1p neurons carrying a LL oscillator have been postulated ^27,66^ and could be a good candidate for adapting evening activity to summer days.

Restriction of the functional clock to the *Clk4.1-GAL4-*expressing DN1p neurons failed to generate LL rhythms (Fig. 3A). However, we observed that PER rescue in most CRY-negative dorsal clock neurons (defined by the *tim-GAL4.cry-GAL80* driver combination) resulted in robust LL rhythms (Fig. 3A). To narrow down the key neuronal group, we tested several sparser *GAL4* lines driving expression in various subsets of clock neurons and observed rhythmic LL behavior when rescuing PER with the *51H05-GAL4* driver (Fig. 3A). The *51H05-GAL4* targets a subset of the glutamatergic DN1ps (Fig. 3B) ^67^. In each brain hemisphere, we observed about 8-9 *R51H05-* labeled neurons, with more than half of them being weakly co-labeled by the *Clk4.1* driver (Fig. 3B-C). The 2-4 *51H05* neurons that were excluded from *Clk4.1* labeling did not express the arousal-promoting ^67^ DH31 neuropeptide (Fig. 3C). In comparison to the *Clk4.*1 DN1ps, the *51H05* DN1p*s* extended additional dorsal neurites in the *Pars intercerebralis* (PI) region (Fig. 3C). Significant non-overlap in the projection patterns of *51H05* and the previously characterized *18H11* DN1p*s* ^67–70^ was also apparent (Fig. 3D). Unlike *18H11*, axons of the *51H05* DN1p*s* scantily innervated the AOTU neuropil, if at all (Fig. 3D). Multi-color flip-out labeling (Fig. 3D) demonstrated that most *51H05* DN1p*s* belongs to the ventrally and contralaterally projecting *vc*-DN1p group ^65,70^, whose role in clock-regulated behaviors remains unexplored ^69–71^.

**Figure 3.**
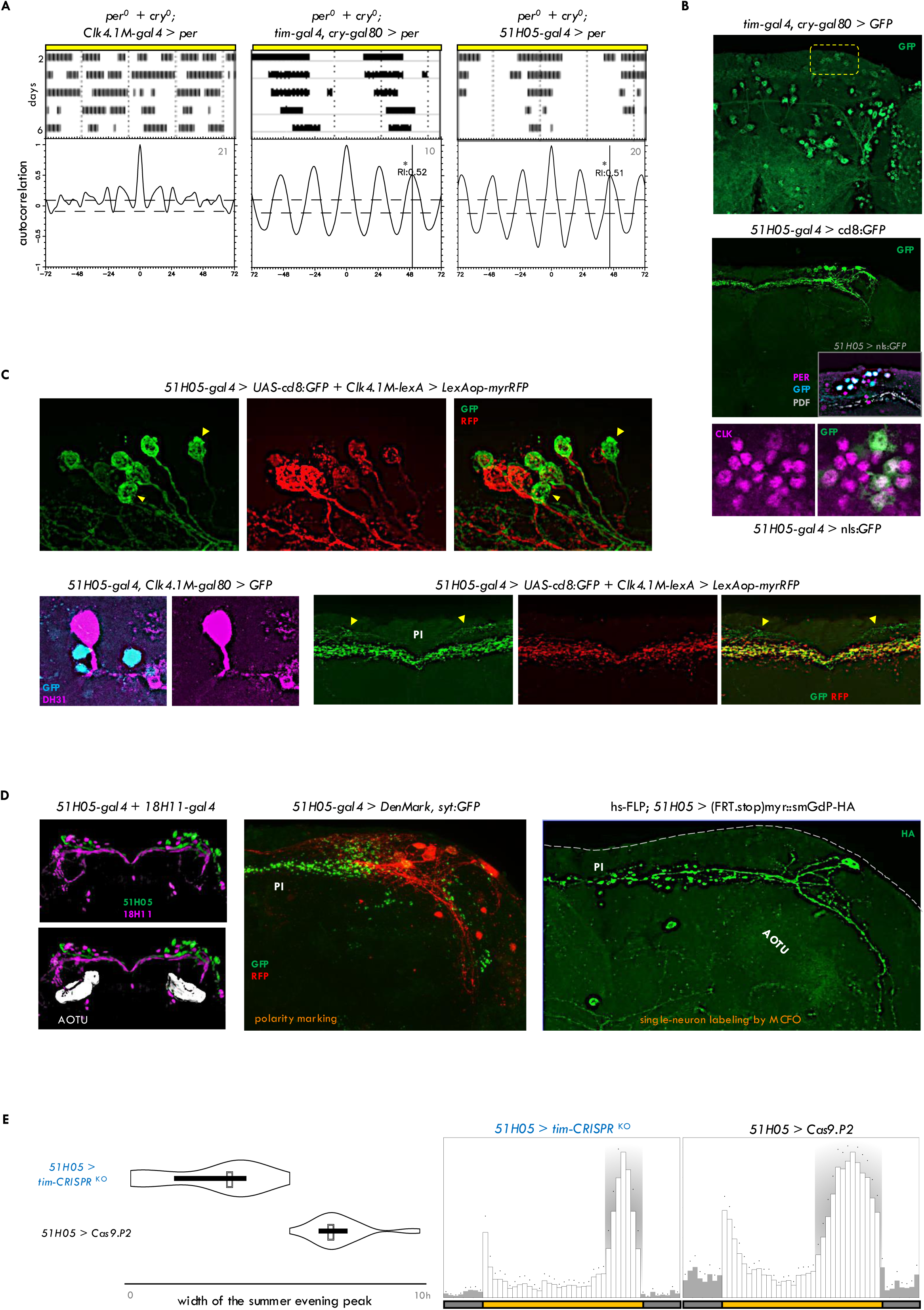
A novel, self-sustaining oscillator in the *vc*DN1p neurons is required for seasonal widening of the evening peak. **A**. Double-plotted actograms of locomotor activity in LL (constant light). In absence of CRY (cryptochrome), functional clock was reinstated in *Clk4.1M-gal4*, *tim-gal4.cry-gal80* and *51H05-gal4* neurons. Autocorrelation analysis shows that both tim-gal4.cry-gal80 and 51H05-gal4 neurons could drive free-running LL rhythms whereas Clk4.1M-gal4 neurons could not. n is marked on the corresponding autocorrelograms. **B**. (Top) Expression pattern driven by *tim-gal4.cry-gal80* (green): the dorsal neurons are highlighted by the yellow box. (Bottom) Expression pattern of *51H05-gal4* (green): 8-9 clock neurons (purple: PER+, CLK+) are detected, belonging to the DN1p cluster. **C**. (Top) Some *51H05-gal4* DN1p neurons (green) are not addressable by *Clk4.1M-lexA* (red), examples are marked with yellow arrowheads. (Bottom left) These *51H05*-positive but *Clk4.1M*-negative DN1ps (cyan) do not express DH31 (purple). (Bottom right) *51H05* DN1ps (green) project additional neurites (yellow arrowheads) to the PI/PL region, compared to *4.1M-lexA* (red). **D**. (Left) While *18H11* DN1ps (purple), as evidenced before, innervate the AOTU neuropil prominently, *51H05* DN1ps (green) barely does. Registration and comparison of Gal4 images on a common template (*Gif*) was through BrainGazer. (Middle) *51H05* neurons send their axons mostly to the PI/PL region, marked by *syt:GFP* (green), and extend ventral dendrites (DenMark: red). (Right) single-neuron labeling by Multi-Color Flip-Out (MCFO) show a 51H05 DN1ps projecting contralaterally and ventrally. **E**. A functional clock in *51H05* DN1ps is required for building the summer-specific expanded evening (E) peak. Expression of *tim*-CRISPR^KO^ in *51H05* collapsed the wide E peak of summer days (p < 0.001 by Mann Whitney U test).

To reveal the clock function of this novel DN1p oscillator, we rendered it clockless by expressing *tim*-CRISPR^KO^ and *cyc*^DN^. Targeted knockout of the clock in *51H05* as well as *tim-GAL4.cry-GAL80* neurons suppressed the broadening of the evening activity in summer-like conditions (Fig. 3E, Fig. S3A). This suggested that the circadian oscillator of the *51H05* DN1ps acts in the evening to either promote a rest-inhibiting function or suppress a rest-promoting function, thereby carving the characteristic evening peak of temperate summer days.

### Working through SIF*a* neurons, *51H05* DN1p*s* link PDF input to daytime rest

To examine the direct impact of the *51H05* neurons on locomotor behavior in summer conditions, we manipulated their firing activity. Silencing *51H05* neurons with Kir2.1 expression ^72^ allowed additional expansion of the summer days’ evening peak whereas hyperexciting them with heat-activated TrpA1 ^73^ constricted it (Fig. S4A). As their constant hyperexcitation recapitulated the phenotype of clock-loss (Fig 3E), we inferred that oscillator function in the *51H05*-DN1p neurons is required for timed downregulation of their rest-promoting neural activity.

To further characterize *51H05* neurons’ function, we activated them in LD 12:12 conditions. Hyperexcitation of the *51H05* (or *tim-GAL4.cry-GAL80*) neurons with TrpA1 abolished the evening peak in equinox days (Fig. 4A). Collapse of the evening peak was still apparent when the overexcitation was restricted to the daytime, or even just the afternoon (Fig. 4A). Optogenetic activation with ChR2-XXL ^74^ underscored that the new DN1p-based oscillator, in stark contrast to the LN-EO, suppresses locomotor activity (Fig. 4B). Video recording of flies undergoing CsChrimson-mediated brief activation of the *51H05* neurons confirmed their lasting quiescence-triggering role (Fig. 4B).

**Figure 4.**
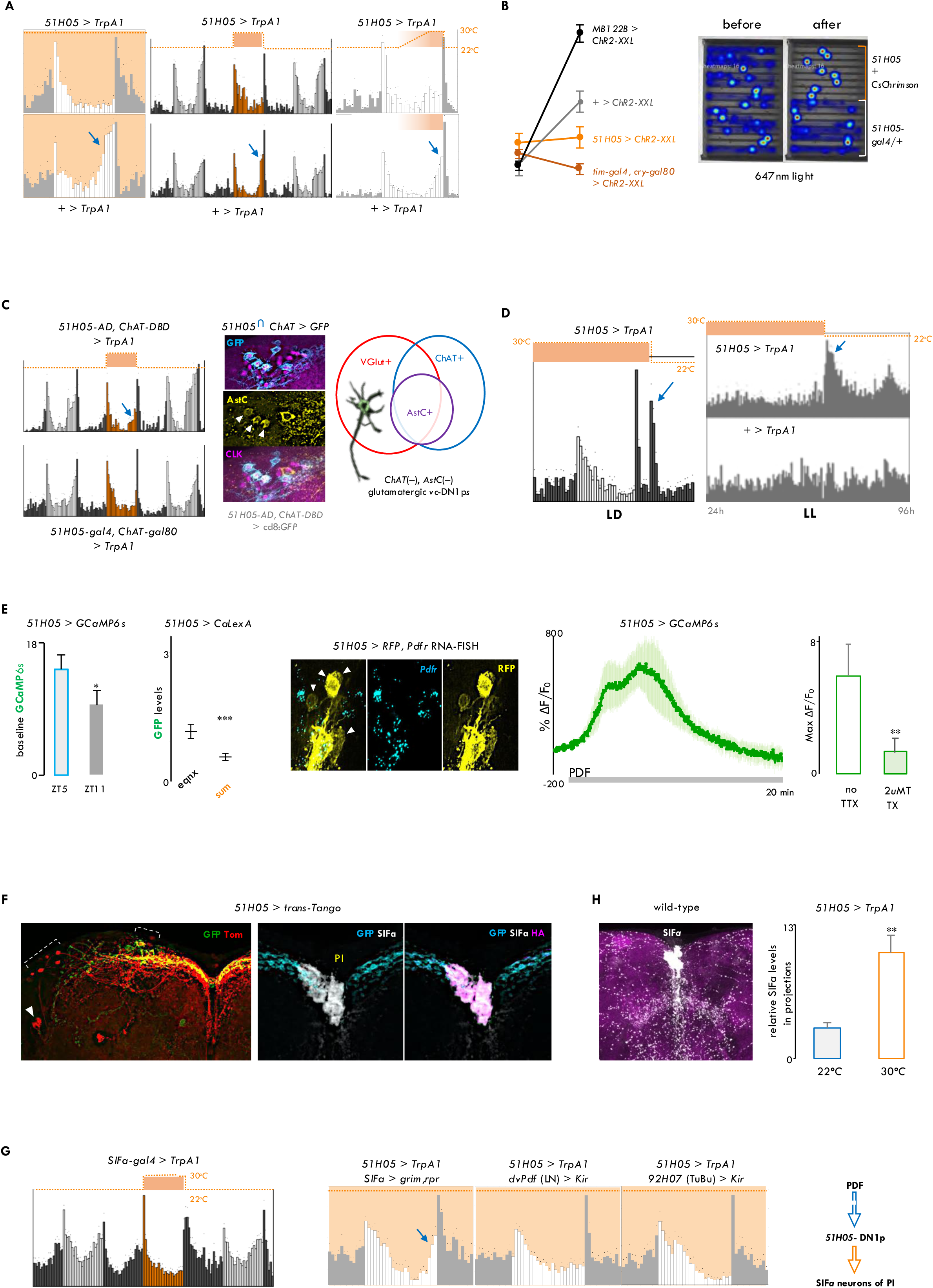
On equinox days, high levels of PDF activate 51H05✍SIF*a* circuit to promote daytime quiescence. **A**. Thermogenetic excitation via expression of TrpA1 channels in the *51H05* DN1ps in equinox condition. (left) Activation by raising the temperature from 22°C to 30°C for 24 hours, (middle) activation during the 12 hour photoperiod, as well as (right) gradually increasing excitation by 1°C step-wise temperature rise in the afternoon suppressed the evening peak (n = 10-16 flies per condition). **B**. (Left) *ChR2-XXL*-mediated optogenetic activation of the DN1ps addressed by *tim-gal4.cry-gal80* and *51H05-gal4* drivers suppressed locomotor activity, as opposed to the arousal-promoting effect of the LN-EO (labelled by MB122B). Average activity in 1hr bin before and during optogenetic activation is shown (n=10-16 flies per genotype, error bars = s.e.m.) (Right) Video recording revealed that *CsChrimson*-mediated brief optogenetic activation of *51H05* neurons rendered flies quiescent. 647nm red light at 4500lux intensity and 5Hz pulse frequency was applied for 10 minutes, fly positions were averaged for 10 minutes (n=8). **C**. (Left) Intersectional drivers showed that overactivation of the non-cholinergic 51H05 neurons (bottom actogram) abolished the evening peak, and not the cholinergic *51H05* neurons (top actogram). n=10-16 flies. (Middle) The AstC neuropeptide (yellow) was restricted to the cholinergic 51H05 neurons (cyan), highlighted with white arrowheads. (Right) Summary of the neurochemical characterization of a novel locomotor-suppressive DN1p subpopulation. **D**. Generation of locomotor activity peaks at night under LD cycles (left), or in arrhythmic LL conditions (right), right after thermogenetic hyperexcitation of *51H05* was lifted by dropping the temperature to 22^0^C. n = 10-16 flies. **E**. (Left) Basal Ca^2+^ activity measured via GCaMP6s reporter suggested that *51H05* DN1ps are more active around midday (ZT5) than toward the evening (ZT11) in LD12:12 (p < 0.05 by *t*-test, n = 8 brains per timepoint). Transcription-based CaLexA-GFP reporter indicated that at midday *51H05* DN1ps are less active under summer conditions (p < 0.01 by Mann Whitney U test, n = 6-8 brains per condition). (Middle) *Pdfr* RNA (cyan) was not detectable in most *51H05* DN1ps (yellow) via RNAscope. Examples of *51H05* soma lacking *Pdfr* expression are highlighted by white arrowheads. (Right) GCaMP6s fluorescence in DN1ps following bath application of 30 uM PDF. The green trace depicts the average of 8 representative neurons’ response from 5 different brains, recorded at ZT6. Preincubation with 2uM TTX significantly diminished PDF-evoked calcium rise (p < 0.01 by *t*-test, n ≥ 5 brains per treatment). **F**. Trans-Tango uncovered LNds (white arrowhead), DNs as well as SIF*amide* expressing PI neurons as direct downstream targets (red and magenta) of the *51H05* DN1ps (green and cyan). **G**. (Left) Overactivation of the SIF*a* neurons for 12 hours during the photoperiod abolished evening peak. (Middle) Functional removal of SIF*a* neurons but not the *dvPdf*-addressed LNs nor the *92H07*-labeled TuBu neurons could rescue the evening-suppressive effect of *51H05* hyperexcitation. (Right) These results from the behavioral epistasis experiments assign the downstream SIF*a* neurons as part of the same evening-suppressive circuit headed by the *51H05* DN1ps, that measures seasonally varying PDF signal. n = 10-16 flies. **H**. 12-hour long overactivation of 51H05 neurons elevated SIF*a* peptide levels in the projections of the PI neurons underscoring functional connectivity (p < 0.01 by *t*-test, n ≥ 5 brains per condition; error bar = s.e.m.).

Taking advantage of intersectional genetics, we proceeded to ascribe the quiescence-promoting role to more restricted subsets of the *51H05* neurons. Hyperexcitation of the *ChAT*(+) *51H05*-DN1p*s* could not abolish the evening peak, while the *ChAT*(–) *51H05*-DN1p*s* could (Fig. 4C). Our results highlighted the non-cholinergic *51H05*-DN1p*s* that do not express the AstC neuropeptide as the core subset of the locomotor-suppressive DN1p*s* (Fig. 4C).

The locomotor-suppressive *51H05* oscillator working alone could produce regular peaks of behavioral activity in LL in the absence of *cry* (Fig 3A). We asked whether timed inhibition of these quiescence-promoting neurons, all by itself, could generate peaks of locomotor activity. Cessation of hyperexcitation of the *51H05* neurons could produce an additional, *i.e.*, third activity peak at nighttime in LD conditions (Fig 4D). The same manipulation could carve an activity peak even when the flies had previously been made arrhythmic by exposure to LL in the presence of *cry* (Fig 4D). Thus, falling neural activity in activity-suppressive neurons can sculpt a peak of locomotor activity.

These results prompted us to posit that the *51H05* DN1p neurons would promote mid-day siesta and counteract the arousal-promotive effect of the LN-EO in the early afternoon. In other words, neural activity of the *51H05* neurons should be high around mid-day and then go down during the afternoon, allowing the manifestation of the evening anticipation. Indeed, basal GcaMP6*s* signal in the *51H05* neurons was significantly lower (Fig. 4E) prior to the lights-OFF transition, when the neural activity of the LN-EO crests ^18,71,75^. On temperate summer days, the evening bout of locomotor activity gets wider and the *51H05* neurons stayed hypoactive as reflected by the CaLexA reporter (Fig. 4E). We thus inferred that the summertime hypoactivity of the *51H05* neurons results from the attenuation of PDF signaling since PDF increased neural activity of the *51H05*-DN1p*s* (Fig. 4E). The lack of *Pdfr* expression in most *51H05* cells (Fig. 4E) suggested an indirect pathway. Indeed, PDF response of these DN1p neurons was downregulated by tetrodotoxin (TTX) pre-treatment (Fig. 4E), corroborating the participation of interneurons in the relay of the PDF message. Notably, *sc*-RNA-based transcriptomic database of the clock neurons ^64^ supports the existence of a glutamatergic DN1p subclass that neither expresses *ChAT*, nor *Dh31*, *AstC* and *Pdfr*.

To explore the output pathways downstream of the *51H05* DN1ps, we employed trans-Tango ^76^. In addition to few DN and LNd neurons, four median neurosecretory cells of the PI were found to be postsynaptic to the *51H05*-DN1p*s* (Fig. 4F). These PI neurons were marked by SIF*amide* expression (Fig. 4F). Notably, a different subset of the *Pdfr*(+) CRY(+) DN1ps were shown to act upstream of the SIF*a* neurons in a memory extinction circuit ^77^. Although SIF*a* has a limited impact on rest-activity rhythms ^78,79^, depletion of SIF*a* leads to hyperactivity ^80^, and activation of SIF*a* neurons promotes sleep ^81^. TrpA1-mediated activation of the SIF*a* neurons during the 12 hours of daytime completely suppressed the evening bout of locomotor activity (Fig. 4G), mirroring the effect of *51H05* neuron overactivation. In fact, ablation of the SIF*a* neurons abolished the locomotor-suppressive effect of *51H05* overactivation (Fig.4G). In contrast, silencing of neither the LNs nor the TuBu neurons of the AOTU, the known downstream targets of the DN1p*s* ^65,68–71,82^, could override the effect of *51H05* activation (Fig. 4G).

Drosophila EM connectome revealed synapses between the DN1p and SIF*a* neurons ^83^. 12-hour-long activation of the *51H05* neurons elevated SIF*a* levels in the projections of the PI neurons (Fig. 4H). Pharmacogenetic excitation of the *51H05* cells induced calcium rise in SIF*a* neurons (Fig. S4C). Taken together, the results indicate that *51H05* neurons directly activate rest-promoting SIF*a* neurons of the PI to attain quiescence. Their reduced activity in low PDF summer-like conditions is thus expected to induce a broadening of the evening peak.

### PDF-induced changes in oscillator coupling confer a prominent role to the DN1p clock in summer-like conditions

We set out to determine how the novel DN1p oscillator senses PDF and imparts its command to the downstream circuit elements. With the SIF*a* neurons of the PI, *51H05*-DN1p*s* displayed prominent GRASP, indicative of their close anatomical proximity (Fig. 5A). Striking diminution of this GRASP signal was seen in summer-like conditions (Fig. 5A). Likewise, diminished GRASP was apparent in flies lacking *Pdfr* (Fig. 5A). Importantly, levels of the SIF*a* peptide in projections of the PI neurons also went down on summer days (Fig. 5B), and *Pdfr*^han^ mutants recapitulated this loss of SIF*a* peptide on standard equinox days (Fig. 5B). Therefore, the feedforward input from the *51H05*-DN1p*s* to the SIF*a* neurons wanes under temperate summer conditions in a PDF-dependent manner, reducing the levels of the SIFa neuropeptide in the brain. We infer that decreased PDF levels in summer conditions would be sensed by the LN-EO to inform the PDFR(-) *51H05* DN1ps about the environment. Notably, *51H05*–LN-EO anatomical connectivity did not change with seasons (Fig. S5A), suggesting targeted structural plasticity in the *51H05–*SIF*a* pathway.

**Figure 5.**
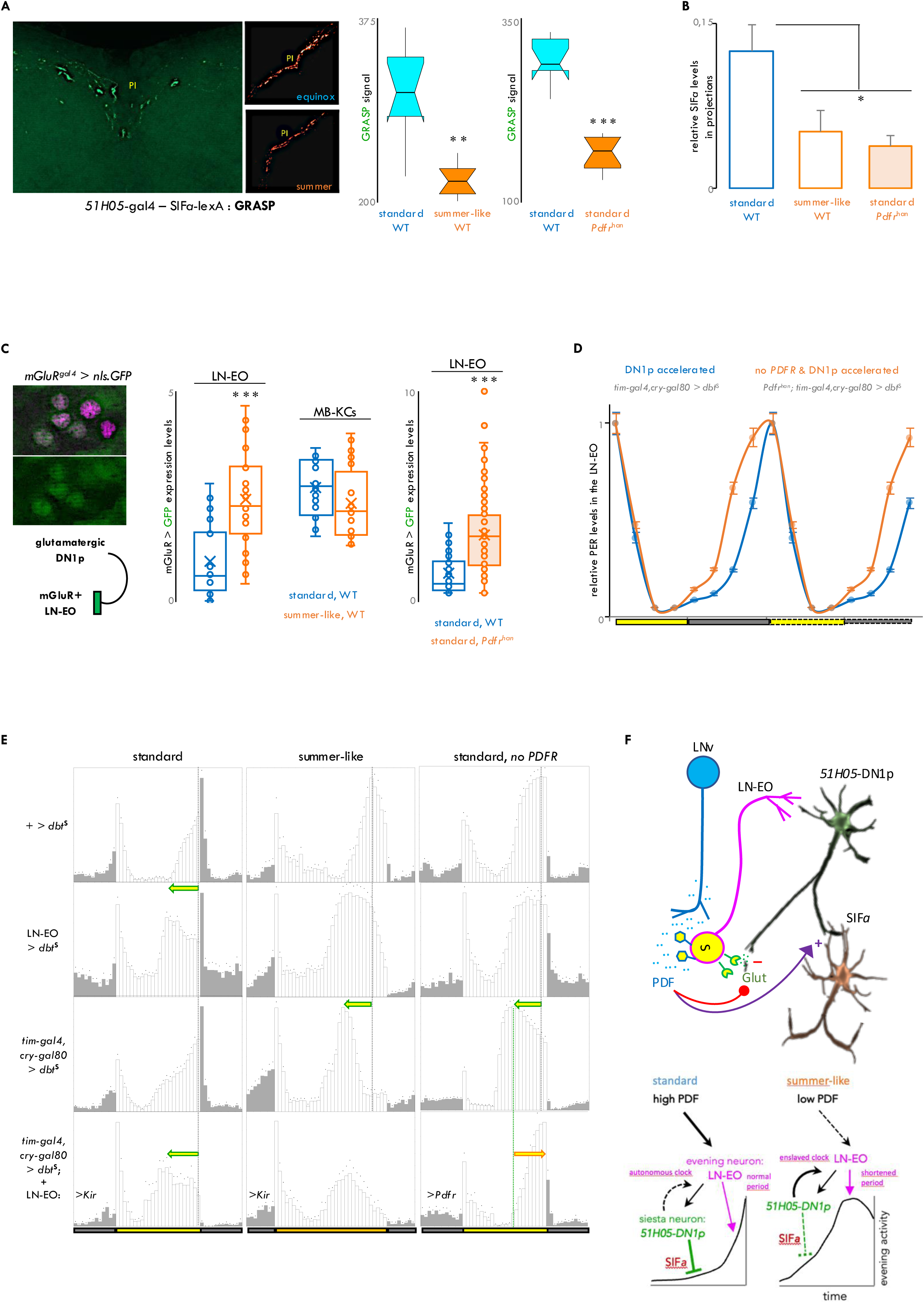
Rewired summer couplings accentuate DN1ps’ status in network hierarchy. **A**. *51H05* DN1ps are in close anatomical proximity of the SIF*a*+ PI neurons; their structural connectivity, as revealed by GRASP, is weakened in summer, or in the absence of *Pdfr* in equinox days (**p < 0.01 and ***p < 0.001 by Mann Whitney U-test, n =6-8 brains per condition, dissected at the middle of the photoperiod). **B**. SIF*a* peptide levels in PI neurons’ projections were diminished in summer, or in the absence of *Pdfr* in equinox days (p < 0.05 by one-way ANOVA with Tukey’s post hoc test, n =5-6 brains per condition, dissected at the middle of the photoperiod). **C**. A T2A-gal4 knockin in the *mGluR* gene drove expression of nuclear-GFP (green) lighting up the LN-EO (PER: purple). DN1 to LNd communication relies on *mGluR* ^68^. Specifically, in the LN-EO neurons, *mGluR* gene expression is upregulated on summer days, or in the absence of *Pdfr* in equinox days. Other *mGluR*-expressing brain regions, *viz*., the Kenyon cells (KCs) of the mushroom body (MB) did not exhibit seasonal plasticity in *mGluR* expression. (***p < 0.001 by Mann Whitney U-test, n = 6-8 brains per condition, dissected at the middle of the photoperiod). **D**. In the absence of *Pdfr*, when there is more mGluR in the LN-EO, acceleration of the DN1p oscillator by DBT^s^ expression resulted in advancement of PER oscillations in the LN-EO, hinting at a strengthened DN1p/LN-EO interoscillator coupling (p < 0.001 by two-way ANOVA, n = 5-8 brains per timepoint, error bars = s.e.m.). **E**. (First-third rows) While in the equinox days, LN-EO clock set the peak phase of evening (E) activity, it is the DN1p oscillator that timed the evening peak on summer days. E phase determination came under the purview of the DN1p clock in equinox days if *Pdfr* was absent. (Fourth row) In equinox days, in the presence of *Pdfr*, phase control could be handed over to the DN1p if the LN-EO was physiologically removed from the circuit by *Kir* expression, revealing conditional hierarchy and unmasking the presence of a LN-EO-independent evening output circuit. *Pdfr* signaling in the LN-EO forms a central hub for organizing the context-dependent hierarchy. DN1ps lost control of phasing of the E-peak in equinox when *Pdfr* was specifically reinstated on the LN-EO. n = 10-16 flies. **F**. Our working model explains how seasonal remodeling of the evening activity bout is accomplished. PDF(+) LNv neurons send input to the *51H05* DN1ps via PDFR-expressing LN-EO neurons. These DN1ps in turn feedback on the LN-EO, and also communicate with downstream SIF*a*(+) medial neurosecretory cells that promote daytime quiescence. PDFR signaling downregulates the DN1p to LN-EO connection and upregulates DN1p to SIF*a* connectivity. On summer days, attenuation of the PDF-PDFR signaling tone weakens DN1ps’ quiescence-promoting output on SIFa neurons and thereby expands the evening bout of locomotor activity. Moreover, attenuated PDF*ergic* tone in summer potentiates the *51H05* DN1p oscillator’s authority over the LN-EO pacemaker, leading to the emergence of a new chain of command for phasing evening activity. The evening oscillators accelerate as PDF levels drop. As a result, the summer evening peak starts to form earlier during the afternoon. Taken together, opposing and flexible changes along distributed circuit nodes orchestrate seasonal expansion and advancement of the evening activity in Drosophila.

*51H05*-DN1p*s* extend prominent projections ventrally toward the LN-EO, the post-synaptic targets of the *51H05* neuron*s* confirmed by *trans*-Tango (Fig 4F). mGluR in the LN-EO mediates glutamate reception from the DN1p*s* ^68^. We asked if this glutamatergic pathway manifests seasonal plasticity. In wild-type flies, *mGluR*-driven GFP levels, a proxy for *mGluR* expression in the LN-EO, increased under summer-like conditions (Fig. 5C). Remarkably, other mGluR(+) neurons, *e.g.*, the Kenyon cells of the mushroom body did not exhibit summer-driven enhancement of *mGluR* expression (Fig. 5C). *mGluR* expression in the LN-EO also increased in *Pdfr*^han^ flies on standard equinox days (Fig. 5C). These results suggested that the connection strength between the glutamatergic DN1p*s* and the LN-EO is increased by weakened PDFR signaling under temperate summer conditions. Therefore, coupling between DN1ps and LN-EO should become stronger in *Pdfr*^han^ flies even under standard equinox conditions. Indeed, when DN1p*s* were forced to run faster, PER oscillations in the LN-EO were advanced in *Pdfr*^han^ mutants in comparison to control flies (Fig. 5D). Summer-like conditions thus increase the control of the LN-EO by the *51H05* DN1p*s*.

To understand how the LN-EO and the *51H05* DN1ps interact to adapt the evening activity to summer-like conditions, we manipulated the clock of each oscillator and assessed the behavioral consequence in standard and summer-like conditions, as well as in standard conditions with no PDFR signaling. Accelerating the ITP+ clock neurons (key constituent of the LN-EO) advanced the evening peak more in standard conditions than in summer-like conditions (Fig. 5E, Fig S5B). In contrast, acceleration of the DN1p clock had no effect in equinox conditions but strongly advanced the evening peak on summer conditions or in the absence of PDFR (Fig. 5E, Fig S5B). Thus, absent or reduced PDFR signaling allows DN1p to conditionally gain control over the phasing of evening activity, whereas high PDF cancels their contribution to phasing. Selectively restoring PDFR signaling in the LN-EO rescinded the control of the DN1p*s* on evening phase (Fig. 5E, Fig S5B). Hence, PDF signal is measured by the LN-EO which then gates the contribution of the DN1p clock. When the LN-EO was incapacitated through *Kir* expression, the phase of the evening peak was determined by the DN1p clock in both standard and summer-like conditions (Fig. 5E). Altogether, our data support a model where PDF sensing by the LN-EO, on one hand, increases the rest-promoting function of the 51H05 DN1ps, and on the other hand, decreases the contribution of the *51H05* DN1p clock to the phasing of the evening peak (Fig. 5F).

## Discussion

Adapting the activity profile to seasonally varying light and temperature conditions is a key part of fitness in non-equatorial animal species. Our work uncovers the molecular and circuit bases of this adaptive plasticity in Drosophila.

### Neuropeptide dynamics mirror external environmental conditions in the brain

Our behavioral data reveal that flies exposed to temperate summer days show a broader and advanced evening activity in comparison to standard laboratory conditions (equinox days). This behavioral adaptability is largely reduced in flies lacking PDF, suggesting that PDF neuropeptide is involved in adaptation to summer days. Indeed, wild-type flies have decreased levels of both PDF and PDFR in temperate summer days. The advanced evening peak and the decreased PDF levels in summer conditions are also observed in the *D. mercatorum* and D. *malerkotliana* species, which have a broad latitudinal distribution. Since these correlated behavioral and PDF changes are not recapitulated by equatorial drosophila species like the *D. Yakuba*, we hypothesize that PDF-dependent behavioral change is restricted to species that encounter strong seasonal variations in their environment. Indeed, in contrast to the *D. melanogaster* flies, photoperiodic adjustment of the evening peak is also absent in the equatorial island species *D. Sechellia*, and this difference has been attributed to divergence in *Pdf* gene regulation between these two species^84^. At the other extreme, the evening peak of many subarctic *Drosophila* species under polar summer conditions also shows greater prominence, marked by a broader activity window and an advanced phase ^7,35^. In the temperate zone, distinctly long daylength occurs around the summer solstice, yet daily temperature and light maxima rarely reach obnoxious levels. Over the late afternoon, weather conditions in the temperate zone likely become optimized for the heterothermic fruitfly to be most active at least energetic cost. Indeed, melanogaster flies under lab-simulated as well as outdoor experiments tested in Leicester (52.6°N) around mid-July displayed mostly a single prominent daily activity peak in the late afternoon, shaped broadly and advanced in phase ^31,85^. Our findings indicate that PDF plays a major role in this adaptation to long photoperiod, low light, and warm temperatures, characteristic of the temperate zone summer days.

The decrease of PDF immunoreactivity in temperate summer days is associated with lower *Pdf* mRNA levels, particularly in sLNvs, suggesting a transcriptional control. Under LD laboratory conditions, PDF immunoreactivity cycles ^86^ but *Pdf* mRNA does not ^86,87^. On the other hand, high light induces *Pdf* mRNA increase in the sLNvs ^10^, while sub-physiological cold temperatures ^88^, characteristic of harsh winter days, decrease *Pdf* mRNA ^38^. Collectively, these findings highlight that the regulation of *Pdf* mRNA levels is crucial to environmental context-dependent PDF dynamics.

Wild-type flies tend to build their evening activity earlier in the afternoon at moderately low temperatures ^5^. The previously observed decrease in PDF levels in cold winter days could thus contribute to this phase advancement. Conversely, under long photoperiod (LD 16:8), wild-type melanogaster flies delay their evening activity peak, while flies lacking PDF cannot ^43,48^. Remarkably, even a drastic change in photoperiod from 8 to 16 hours does not significantly alter the mean PDF levels or the amplitude of PDF cycling in the sLNvs ^38^. Therefore, the marked decrease in PDF observed during temperate summer days is likely due to synergistic interaction between low light intensity, moderately high temperature, and long photoperiod.

PDF thus appears to be a key component to modify the locomotor behavior according to environmental variations, thus allowing adaptation to daily or seasonal environmental changes in temperature, light intensity, and photoperiod. Being controlled by light, temperature as well as the circadian clock, PDF levels are logically central in phasing activity within the 24h day-night cycle.

### PDF Guides Phase Plasticity via an Inhibitory PKA–SGG Pathway Targeting TIM

How does the biochemical clock program get reset by PDF still remains unclear. PDFR signaling, particularly through the cAMP/PKA pathway, post-translationally controls PER/TIM half-life but the exact effects seem to be diverse and cell-specific ^57,58,89^. In addition, PDFR-dependent transcriptional regulation of clock genes affects the amplitude of the molecular rhythms ^59,90^.

Our results show that the LNv-derived PDF decelerates the PDFR+ LNd oscillator, highlighting a causal basis of the PDF-induced delay in the phase of calcium oscillations and behavioral output generated by the LNd’s evening oscillator ^42,75^. Subsequently, we implicate GSK3/SGG as the core clock component directly translating PDF-evoked upsurge of PKA activity into pace change of the LNd oscillator. Our data strongly suggest that SGG is part of the PDFR signaling pathway and is a direct target of PKA-catalyzed inhibitory phosphorylation. In the *Drosophila* clockwork, the principal substrate of SGG kinase is TIM ^60,61^. On the S297 and T301 residue, SGG-catalyzed TIM phosphorylation promotes its nuclear accumulation, albeit the strength of this effect widely varies across clock neurons ^61^. Our work hints that these phosphorylation events facilitate additional SGG-regulated TIM modification, resulting in increased TIM turnover. Notably, sequential and hierarchical phosphorylation events are also hallmarks of CK1/DBT’s catalysis on PER ^91^. PDFR and SGG-influenced regulation of TIM stability and oscillations was potent in the oscillator of the key ITP+ evening neurons and not on the PDF+ morning neurons (sLNv) that also express PDFR. The distinct composition of the PDFR signalosome between morning and evening neurons ^92^, and the distinct architecture of the biochemical clock program itself ^93^ could underpin the differential effects of PDFR on morning/evening oscillators.

Inhibitory S9 phosphorylation of SGG is strongly regulated by PKB/AKT signaling, and sensitivity of the *Drosophila* behavioral rhythms to nutrient status and energy level is thought to be mediated by AKT and TOR pathways acting on SGG activity to control nuclear accumulation of TIM ^94^. Furthermore, SGG’s action on TIM also interfaces with CRY-dependent TIM-degradation induced by light. Light-responsive serotonin/5HT1B signaling modulates TIM stability through S9 phosphorylation of SGG ^95^. TIM is the central determinant of the clock’s photosensitivity, with *tim* different alleles regulating the adaptation of the behavior to the photoperiod as a function of latitude ^96–100^. Taken together with our results suggesting seasonally varying SGG-TIM interaction, TIM indeed appears to sense a series of internal states and environmental changes to translate these signals into modifications of the molecular clock. The LNd evening neurons may be uniquely able to segregate the effect of different upstream pathways on SGG-TIM interaction by opting for time-dependent usage of alternatively spliced SGG isoforms ^101^.

### Neural Circuit Mechanism Underlying Seasonal Waveform Plasticity

We identified a small subset of glutamatergic DN1p neurons as key regulators of siesta. These 2-4 CRY-negative neurons are best characterized by the expression of the *51H05-Gal4* driver, which can autonomously drive sustained behavioral rhythms in LL. The DN1ps are highly heterogenous ^64^, and are involved in different sleep-promoting and sleep-inhibiting effects through other clock neurons or downstream non-clock cells ^65,67–71,102^. The DH31-negative, CRY-negative *51H05* neurons belong to the anatomically defined *vc*-DN1p group ^70^, whose function remained unknown until now. We demonstrated that a functional clock in these DN1ps is required to broaden evening activity in temperate summer days. Overactivation of these neurons extends the afternoon siesta, whereas silencing them widens the evening peak of activity to the afternoon. Importantly, PDF increases the neuronal activity of the *51H05* DN1ps. Hence, temperate summer days with low PDF levels keep them hypoactive allowing the evening peak to expand into the afternoon.

This new DN1p siesta-promoting circuit appears to be unrelated to the previously described DN1p–TuBu sleep-promoting and sleep-inhibiting circuits ^65,68–71^. Whether PDF has some effects on the other DN1p-controlled circuits is unknown, although PDFR is expressed in the CRY-positive glutamatergic DN1ps ^64^. Despite being PDF-regulated *51H05* DN1ps are PDFR-negative cells, indirectly receiving PDF signal through PDFR-expressing LN-EO that sets the phase of the evening activity in standard equinox days. Our results also reveal a complementary interaction between these two subsets of the evening neurons. While accelerating the *51H05* DN1p clock in standard days does not affect the evening peak of activity, it dramatically advances evening activity by several hours in temperate summer days, or in the absence of PDF signaling. In these summer conditions, a decrease of the sleep-promoting effect of the *51H05* DN1ps is thus associated with an increase of their contribution to the pace of the circadian network.

How could such changes occur? Internal state-dependent, axonal branch-specific anatomical plasticity is shown by specific dopaminergic neurons (PPM2) in the fly brain ^103^. In the clock network, PDF-expressing sLNvs show a daily rhythm of axonal arborization expansion ^104–106^. This arborization is required for the integration of temperature inputs through glutamatergic DN1ps ^107^ and the daily cycling is regulated by PDFR signaling ^108^. It is thus possible that the LNd/DN1p switch and/or the decrease of DN1p activity in low-PDF temperate summer days are regulated by related mechanisms. Similarly, on temperate summer days, *51H05* DN1p neurons alter their medial connections with the SIF*a* neurons of the PI, while maintaining their ventral projections to LN-EO. We hypothesize that in low-PDF summer conditions, the hypoactive glutamatergic DN1ps strengthen their functional connections with LN-EO, through increased postsynaptic mGluR expression and possibly enhanced glutamate release. Differential regulation and release of co-transmitted neurotransmitters/neuropeptides based on firing patterns and rates, as widely seen in both invertebrate and mammalian neurons ^109,110^, supports the potential for selective strengthening of DN1p glutamatergic synapses in their ventral branch under low-PDF conditions. Given that DN1ps, like SCN clock neurons, co-express multiple output molecules, selective and branch-specific synaptic adaptation likely represents a general mechanism for modifying the circadian neural circuitry based on internal and external states.

This switch from the LN-EO clock to the DN1p clock under low PDF conditions may enhance the plasticity of evening activity in response to environmental changes, as the DN1p clock neurons are more sensitive to temperature and light fluctuations ^10,46,52,65,111,112^. On harsher Mediterranean-type summer days, light, temperature, and PDF counteract circadian signals, delaying evening activity closer to nightfall ^48,84^. However, during temperate summer days, the circadian network takes the lead through the otherwise hypoactive *51H05* siesta neurons to, on one hand, keep evening activity elevated in the afternoon, and on the other hand, to accelerate the main evening clock, *i.e.,* LN-EO. We speculate that higher latitude-restricted drosophilids have evening circuits permanently locked in the latter configuration.

### SIFamide Neurons Link the Clock to Seasonal Changes in Systemic Physiology

We show that the locomotor-suppressive dorsal clock neurons communicate with a defined subset of neurosecretory cells in the pars intercerebralis (PI) that express the neuropeptide SIFamide, and that activation of *51H05* neurons induces an increase in SIFa levels. Only 2 pairs of SIFa neurons extend projections onto the entire *Drosophila* CNS to regulate reproduction, food intake, and sleep ^80,113–115^. 51H05 DN1ps need SIFa neurons to promote siesta and overactivated SIFa neurons suppress the evening peak of activity, in agreement with their sleep-promoting function ^80,81^. Under temperate summer conditions or in the absence of PDF signaling, the anatomical connection between DN1ps and SIFa neurons is reduced and SIFa levels decrease. A drop of orexigenic SIFa, commanded by the upstream PDF-DN1p circuit, thus permits the fly to sustain high-level locomotor activity during temperate summer days. Remarkably, a similar circuit mechanism reliant on shifting connectivities of a separate LNv-DN1a circuit also underlies daytime-dependent plasticity of light-evoked locomotor response ^140^.

In mammals, efferent SCN neurons characteristically restrict their innervation within the hypothalamus. The neuroendocrine command center, *i.e.*, pars intercerebralis of the insect brain displays physiological and developmental equivalence with the vertebrate hypothalamus ^116^. The pars intercerebralis houses regulatory centers for sleep, arousal, and rhythmic rest-activity, as does the hypothalamus in mammals ^79,117,118^. Notably, the evolutionary homolog of SIFaR in vertebrates NPFFR1/2, senses RF-amide-related peptides (RFRPs), whose hypothalamic expression varies seasonally ^119^. Widespread control of vertebrate seasonal reproduction by RFRP neurons that receive input from the SCN ^119^ suggests a potential role of SIFamide in bridging insect reproductive physiology with the brain circadian network.

While suppressing sleep, a reduction in SIFamide under temperate summer conditions should also facilitate mating at the expense of feeding, and consequently, reduce lifespan ^113,114^. This behavioral goal-switch is complemented by overactivated insulin/dilp2 signaling, driven by the fly’s preference for yeast intake during hot summer days ^120,121^. The clock neuronal network thus uses parallel light-modulated outputs via DN1p–SIFa and LNv–LHLK pathways that integrate sleep with hunger ^122,123^. It will be interesting to see how these circuits coordinate to control siesta and evening activity on summer days as a function of light levels and feeding status.

## Materials and methods

### Fly lines

A list of the fly lines, their source and corresponding references are provided in a supplementary excel table. All fly stains were maintained on corn meal media at 25°C in 12:12 LD.

### Generation of transgenic flies

#### ITP-gal4

To obtain the ITP-gal4 line, the gal4 sequence included in the vector pBS-KS-attB1-2-GT-SA-Gal4-Hsp70pA (DGRC stock# 1325) was inserted in place of the transposon <u>Mi{MIC}ITP</u>(MI00349) contained in the fly stock RRID:BDSC_30713 using Recombination Mediated Cassette Exchange (RMCE) technique ^124^ as previously described ^10^. Transgenesis was performed by BestGene.

#### LexAop-PDFr-mCherry

LexAop-PDFr-mCherry transgene was constructed by replacing the myr-GFP sequence of plasmid pJFRC19-13XLexAop2-IVS-myr::GFP (pJFRC19 addgene Plasmid #26224) with the PDFr-mCherry sequence as described below. The PDFr ORF (from exon2 excepted 27 first base pairs to exon9 before last codon) sequence ends with Glutamine in frame with mCherry sequence. PDFr-mCherry sequence was PCR amplified from c13758 mCherry pcDNA3 (gift from Paul Taghert) using two primers containing respectively SalI and XbaI restriction sites: For 5’AT*GTCGAC*ATGACCCTCCTGTCGAACAT3’ and, Rev 5’CC*TCTAGA*GGCTACTTGTACAGCTCGTCCATG3’. It was then inserted in pJFRC19 previously digested by XhoI and XbaI restriction enzymes using T4 DNA ligase. After confirmation by sequencing, this construct was introduced inside the 3rd chromosome attP docking site VK00005 doing a phiC31 integrase-mediated transformation (BestGene).

Transgenes for bimolecular fluorescence complementation (BiFC) technology: Transgenes UAS-pkaC1-VenusC, UAS-pkaR2-VenusC and UAS-VenusN-shaggy were constructed in two steps. Firstly, DNA fragments coding for C-terminal 155-238 and N-terminal 1-154 parts of Venus protein were PCR amplified from respectively plasmids UAStHth-VC and UASTsh-VN (gift from S. Merabet). Primers For 5’*atatCTCGAG*GCCGACAAGCAGAA3’ plus Rev 5’*tgcgTCTAGA*TTACTtgtACAGCTCGTCC 3’ and for 5’att*CTCGAG*ATGGTGAGCAAGGGAGA3’ plus Rev 5’aat*TCTAGA*tccaccactacctccGGTGATATAGACGTTGTGG3’ were used respectively. Those primers include XhoI and XbaI restriction sites to allow the insertion in the new plasmid, and when necessary, a linker of five amino acids GGSGG were added to separate the Venus fragment from the protein of interest, as well as bases to keep the frame with shaggy coding sequence. Each Venus moiety was inserted in a plasmid containing UAS sequence (pJFRC-MUH, addgene 26213) via XhoI and XbaI sites. Secondly, the entire coding sequence of pkaC1, pkaR2 and shaggy were PCR amplified from respectively FMO06689, FMOO13004 and LD44595 (DGRC clones, Indiana University) and consequently cloned using the SLIC method ^125^ into pJFRC-MUH-VenusC for PkaC1 coding sequences and pJFRC-MUH-VenusN for shaggy coding sequence. PkaR2 sequence were cloned into pJFRC-MUH-VenusC using classical cloning technics. Primers For 5’ATACAAGAAGAGAACTCTGAATAGATCTGCGGCCGCCAGTCGACATGGGCAACAAC3’ and Rev5’GCCTTGATGCCGTTCTTCTGCTTGTCGGCCTCGAGTCCACCACTACCTCCGAATTCA 3’ were designed for pkaC1, For 5’*ATTgcggccgc*ATGTCGAGCGATTCGAGTCG3’ and Rev 5’AA*gtcgac*tccaccactacctccCAAATTCGTATTACGGCGG3’ for pkaR2, For 5’ CTATATCACCggaggtagtggtggaTCTAGAcacggtcacgctttacagtc 3’ and Rev5’AAGTAAGGTTCCTTCACAAAGATCCtctagaTTTTTTTTTTTTTTTTTTACT3’ for shaggy. BestGene did the transgenesis using the PhiC31 integrase system. UAS-pkaC1-VenusC was inserted in the attP40 docking site 2L (25C6), UAS-pkaR2-VenusC and UAS-VenusN-shaggy were inserted in the attP2 docking site 3L (68A4).

### Behavioral experiments

All behavioral experiments were performed with adult flies aged between 3-6 days. These flies were raised at 25°C and in 12:12 LD in mixed-sex groups. The activity-rest rhythm was recorded in Drosophila Activity Monitor (DAM2 5mm; TriKinetics) as previously described. For the bigger *D. mercatorum* flies 7mm DAM2 tubes were used. 25°C temperature, 12h photoperiod and 500-1000 lux (555 nm) light intensity were used for all standard laboratory equinox experiments, unless otherwise mentioned (*e.g.*, for TrpA1 activation). Summer-like conditions were imposed by a constant 28°C temperature and 16h photoperiod. Adult *Drosophila melanogaster* flies dwell on fallen plant material in fruit orchards, woodlands, and forests ^126,127^. From boreal to sub-tropical zones, seasonal changes in foliage lead to a dense canopy on summer days and lower light flux at the understorey of fruit orchards and woodlands is a common feature ^128–131^. Consequently, flies are paradoxically exposed to lower light intensity in summer months compared to spring and autumn days ^132–134^. Furthermore, outside the tropics, twilights marked by low-intensity light availability are substantially longer during summer ^135^, strengthening the association of temperate summer twilights with low light for the crepuscular flies. In light of these, we used lower light intensity (50 lux, using gray neutral density filters) for summer-like conditions.

For RD experiments, red light was administered by 620-650nm LEDs (Lunartec, Pearl diffusion). USB200 (Ocean Optics) spectrometer was used to measure light spectra and irradiance. For TrpA1 experiments, temperature of the Percival incubators was changed by software control either in 1°C step/hr or directly and temperature values were crosschecked with independent loggers. Data were analyzed using FaasX software, which is derived from Brandeis Rhythm Package. FaasX is compatible on Apple Macintosh OSX computers and is freely available (see https://neuropsi.cnrs.fr/en/departments/cnn/group-leader-francois-rouyer/). Activity data are presented as an average of 5-6 days and of *n* flies leaving the first 2-3 days of the experiment out. On the average activity profiles, each gray/white bar denotes the mean activity levels in a 0.5h interval during the light phase and each black bar denotes the mean activity levels in a 0.5h interval during the dark phase of the LD cycle. The dot on each bar shows the s.e.m. For the LL experiments, rhythmic flies were defined by autocorrelation (RI jitter = 5 bins and maximum lag = 144 bins) function of FaasX. For optogenetic experiments, 1-2 days old flies were starved overnight and fed 400uM all-trans Retinal (ATR in 5% sucrose and 2% LMP agar) for 3 days before starting the experiment during which they continued to feed on a new batch of ATR-laced food. For video analysis of CsChrimson activation, Noldus suite was employed and EthoVision software was used to produce the location heat maps.

### Immunostainings

The immunolabelings were performed using whole-mounted adult *Drosophila* brain. Briefly, brains were dissected in cold PBS, fixed in 4% PFA for 1h at room temperature, washed 6x in 10min steps with PBST (PBS containing 0.3% TritonX) and permeabilized with PBS containing 1% TritonX before undergoing overnight blocking in 1% BSA solution. Primary antibodies – anti-PER (rabbit; 1:15000), anti-PDF (mouse; 1:20000), anti-GFP (chicken; 1:1000), anti-CLK (guinea-pig; 1:2000), anti-TIM (rat; 1:10000), anti-dsRED (rabbit; 1:2000), anti-S9P-SGG (mouse; 1:20), anti-SGG (mouse; 1:500), anti-SIFa (rabbit; 1:5000), anti-HA (rat; 1:200), anti-AstC (rabbit; 1:250), anti-DH31 (guinea-pig; 1:5000) were diluted in PBST containing 0.1% BSA and used for 2-3 nights at 4°C. After washing brains were treated with secondary antibodies for 2 nights at 4°C, washed and mounted. Secondary antibodies were Alexa350-, Alexa488-, Alexa547-FP546-, FP568-, and Alexa647-conjugated goat antibodies (1:1000-1:2000 dilution) directed against IgGs of the appropriate species (Invitrogen). To test the effect of PDF on TIM stability on live fly brains (pulse-chase experiment), dissection was undertaken in AHL at room temperature, under red light (>600nm plastic filter). Brains were incubated in AHL containing 100 uM PDF peptide and kept in an incubator at dark 3.5 h. Then the brains were washed, fixed and processed as above. Fluorescence signals were analyzed with a Zeiss AxioImager Z1 semi-confocal microscope equipped with an AxioCam MRm digital camera and an ApoTome2 module set with an automatically adjustable grid which provided structured illumination. Fiji software was used to quantify fluorescence intensity from digital images. Integrated densities over a defined thresholded area of the axonal arbors of the s-LNvs acquired with a 40x objective were analyzed for quantifying signal in the dorsal projection of the PDF neurons as well as for quantitative GRASP measurement. Use of the wand tool of Fiji yielded equivalent result. For all quantifications from soma, we utilized the formula: I = 100x(S-B)/B to calculate the fluorescence percentage above background (S = mean intensity inside the cell, B = the mean intensity of the region adjacent to the positive cell). For studying clock protein oscillations, a 63x objective was used to acquire images.

### Proximity Ligation Assay (PLA)

Flies expressing MYC-tagged Sgg10 and GFP-tagged PKA (both regulatory and catalytic subunits were tagged with GFP) pan-clock neuronally by *Clock*^6939^*-Gal4*, were dissected in ice-cold PBS around ZT2-4. Brains were fixed in 4% paraformaldehyde for 1 h at room temperature. The whole mounts were treated for regular immunohistochemistry until the addition of the primary antibodies, α-MYC (Novus 9E10; 1:400; rabbit) and α-GFP (Thermo-Fisher A11120; 1:2000; mouse). After 48 hours, brains were washed and then incubated with the anti-rabbit and anti-mouse secondary antibodies conjugated to PLUS and MINUS PLA probes (see ^136^). T4 Ligation and rolling circle amplification of the probes were undertaken using Duolink^TM^-provided reagents. Brains were mounted in Prolong-Diamond. The appearance of fluorescent spots signifying successful amplification was detected by a Leica SP8 confocal microscope. No fluorescent spots were detected when the fly expressed MYC-tagged sgg10 alone. Moreover, when the entire procedure was performed without adding the probes, no fluorescence was apparent.

### Bimolecular fluorescence complementation (BiFc) assay

BiFC was performed essentially as previously described, with necessary adaptation of the technique for adult brains ^137,138^. Transgenes UAS-pkaC1-VenusC, UAS-pkaR2-VenusC and UAS-VenusN-shaggy were constructed as mentioned above. Binary combinations of UAS lines were driven by a pan-clock Gal4 line (*Clock*^6939^*-Gal4*). Brains from 2-3 days old adult flies were dissected in PBS and after a quick (10 min) fixation in 4% PFA were immediately imaged in a Leica SP8 confocal microscope using a 20x objective. The quick fixation was necessary for preventing rapid photobleaching. Brains were dissected at ZT2 and ZT14. In the beginning, some of the BiFC brains were additionally co-stained with anti-PDF antibody. 515nm laser was used for Venus excitation, while emission was collected around 528nm using LasX software interface. Identical parameters of confocal acquisition were applied for every treatment.

### mRNA In situ hybridization (ISH)

The RNAscope® Multiplex Fluorescent Reagent Kit v2 (ACD Bio) was used on whole mount Drosophila adult brains as previously described ^139^. Dissected brains were fixed, washed and then treated with 3% hydrogen peroxide (3% H_2_O_2_) for 20 min at room temperature (RT). The target retrieval treatment was performed for 2 min at 95 °C in 1× Target Retrieval solution and subsequently the brains were treated at RT with Protease IV for 30 min, and washed. The probes (*Pdf* probe: Cat. No. 457471-C3 and *Pdfr* probe: Cat. No. 576571-C2, created by ACD Bio for the present study) were warmed to 40 °C and added to the brains incubated with the probe diluent solution after 1:50 dilution. After overnight incubation at 40 °C, brains were washed and incubated with 2–3 drops of RNAscope® Multiplex FL v2 at 40 °C for 30 min and rewashed. These steps were repeated with Amp 2 and Amp 3 incubations, followed by C2 and/or C3 treatments. In the last step, the samples were incubated for 30 min at 40 °C with Opal 520 (FP1487001KT, 1:2000 dilution) and/or Opal 650 (FP1496001KT, 1:2000 dilution). The brains were washed and incubated overnight at 4 °C with anti-PDF or anti-RFP antibodies. Brains were imaged with semi-confocal microscope and quantification of signal intensity was carried out as for immunostainings.

### GCaMP imaging

Adult Flies were dissected under ice-cold AHL for PDF bath-application experiments and under ice-cold HL3 for P2X2 experiments was used. The whole brain explants were placed on 42 mm diameter coverslips treated with Poly-D-Lysine and Laminin. Then the preparation was immersed in oxygenated AHL or HL3. Calcium imaging was performed using a Zeiss Axio Examiner D1 upright microscope with Apochromat 40X W NA 1.0 immersion lens. GCaMP6s probe was excited (25ms exposure time) with a Colibri 470 nm LED light source and images were acquired using AxioCam MRm at 0.5-2 Hz sampling rate. 0.5mM ATP (Sigma-Aldrich Chemical) was used to stimulate the P2X2 channel. When used, 30 uM PDF (PolyPeptide) was added after at least a minute of baseline recording. ATP was dissolved in HL3 solution, whereas PDF was dissolved in 0.1% DMSO in AHL, essentially as in Chatterjee, 2018. Pre-incubation with 2uM TTX was performed for 15-20 minutes, fresh 2uM TTX was re-applied immediately prior to image acquisition (see ^123^). To measure baseline GCaMP6s intensity, live brains in room-temperature AHL were imaged immediately after dissection. *xyzt* stacks were acquired for 1 minute and a maximum intensity projection was rendered, which was subsequently treated as a static image. The average fluorescence of all pixels for each time point in a defined ROI was subtracted from the average background fluorescence of an identically size ROI elsewhere in the brain. The resulting pixel fluorescence value for each time point was defined as trace Fb. Changes in fluorescence were computed as %dF/F0 = ((Fb-F0)/F0) x 100, where F0 is defined as the average background-subtracted baseline fluorescence for the 30-60 frames preceding the stimulus application. Fiji (Image J) software was used for processing and quantifying all images. Maximum GCaMP6s fluorescence change values (Max dF/F0) were determined as the maximum percentage change observed for each trace over the entire duration in a given imaging experiment. Maximum values for each treatment and genotype were averaged to calculate the mean maximum change from baseline.

PDF: Pigment-Dispersing Factor.
PDFR: PDF Receptor.
GSK3: Glycogen Synthase Kinase 3.
SGG: Shaggy.
DN1p: Dorsal Neuron 1, posterior.
SIFa: SIFamide neuropeptide.
LNd: dorsal Lateral Neuron.

## Sup figure legends

**Figure S1.**
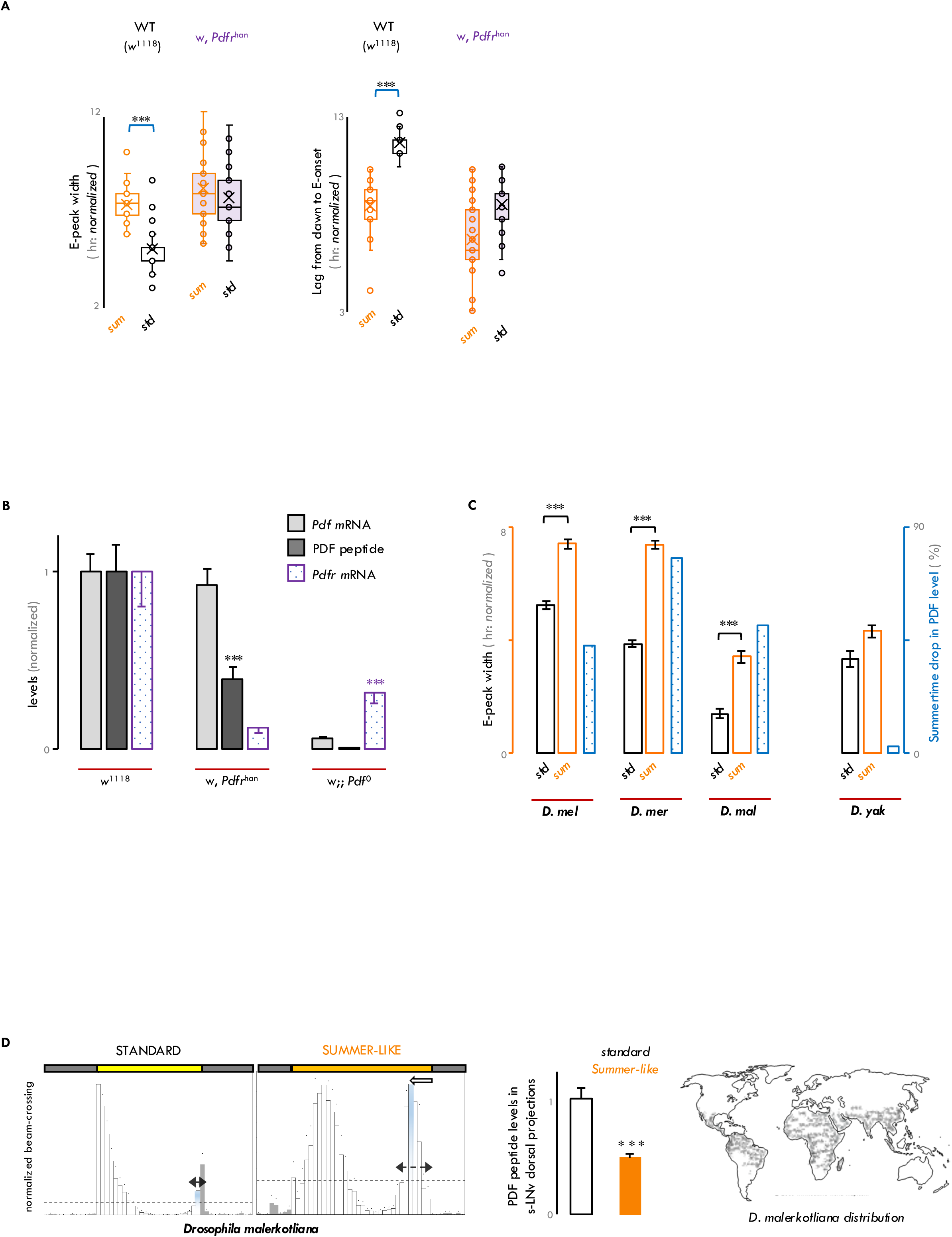
**A**. In summer, the E-activity is both expanded and advanced in wild-type (*w*^1118^) flies. The width of E-peak is broader and the onset of the E-activity is advanced in summer compared to equinox. Both these differences largely disappear in Pdfr^han^ mutants. **B**. The absence of PDFR decreases PDF peptide levels. On the other hand, absence of PDF decreases *Pdfr* mRNA. This is in accordance with our finding that, in summer, not only PDF levels in the LNvs but also its receptor levels drop in the LNd and DN1p. **C**. In summer, broadening of the E-activity is associated with a decrease in PDF levels in different Drosophilids. Cosmopolitan *melanogaster* and *mercatorum* exhibit both broadening of E-activity and drop in PDF levels in summer whereas afrotropical *yakuba* exhibit neither. Moderately distributed *malerkotliana,* like *melanogaster* and *mercatorum* and unlike *yakuba*, exhibit both broadening of E-activity and an associated drop in PDF levels.

**Figure S2.**
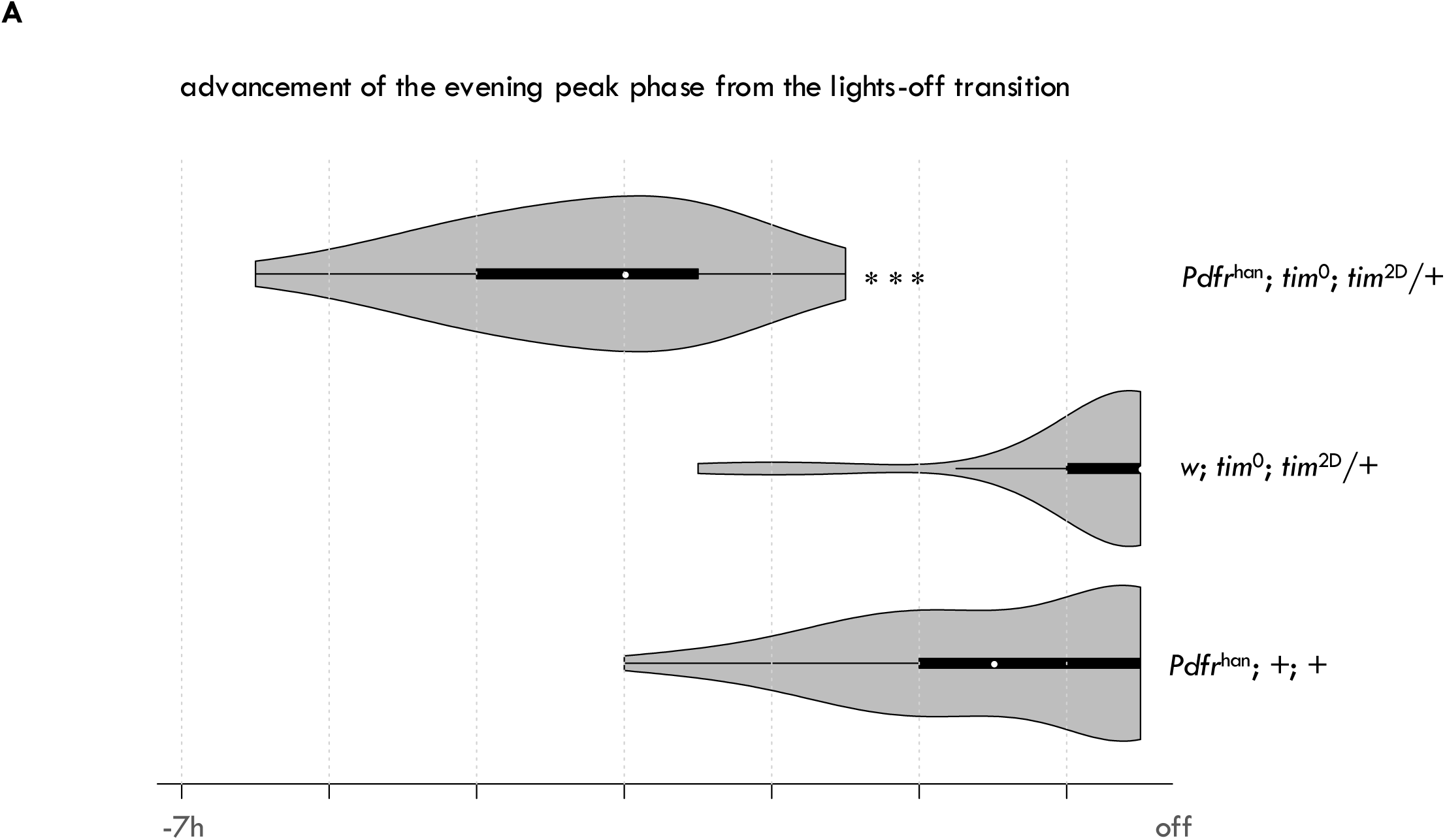
**A**. Quantification of a genetic interaction between a variant of TIM (TIM^2D^) and Pdfr^han^. Advancement of the E-peak phase from the lights OFF transition is greater in the flies with both Pdfr^han^ mutation and TIM^2D^ variant of TIM than flies with only Pdfr^han^ mutation or TIM^2D^ variant of TIM.

**Figure S3.**
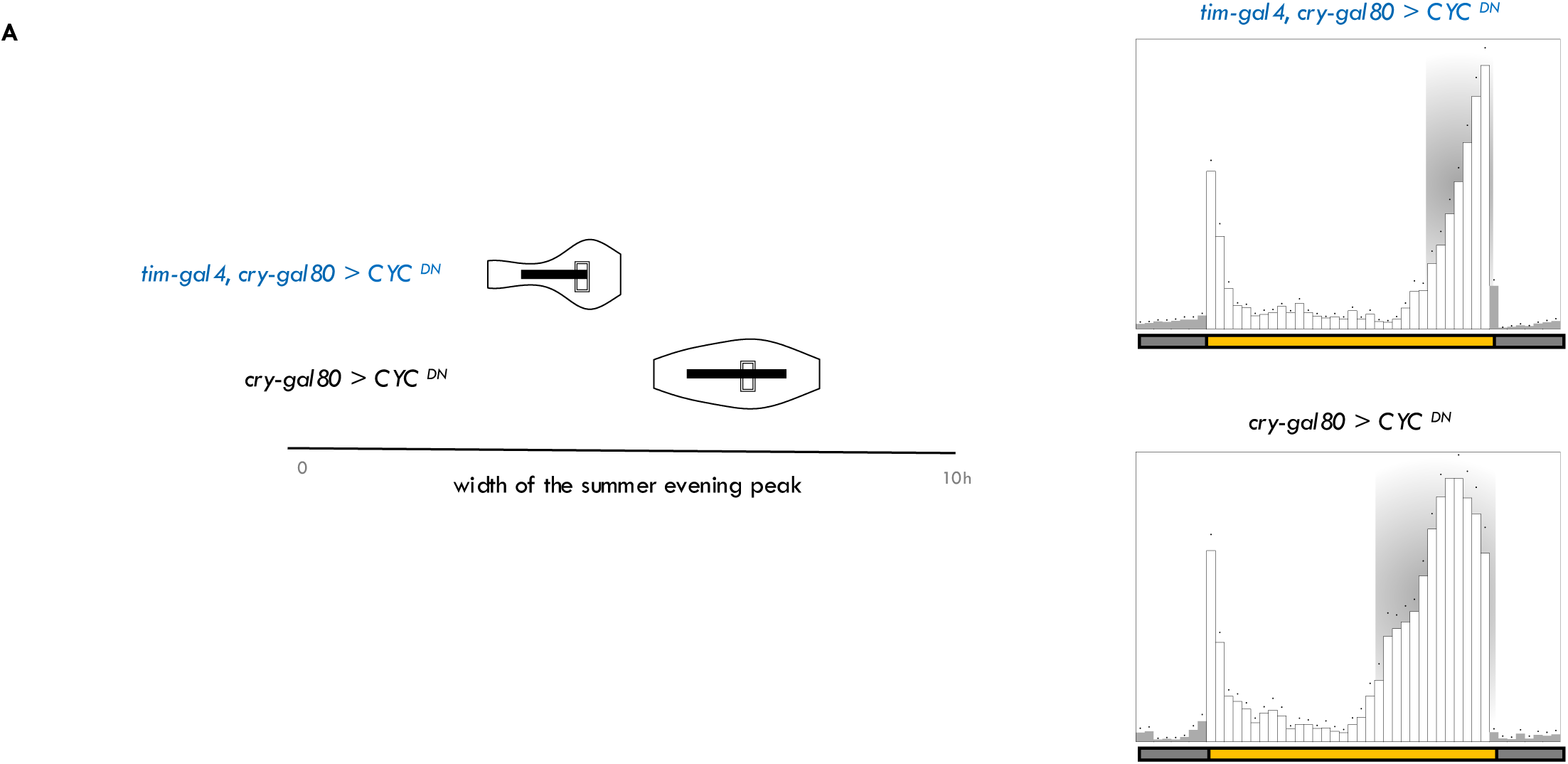
**A**. A functional clock in the tim-gal4, cry-gal80 neurons is necessary for the expanded E-activity in summer. Expression of dominant negative variant of CYCLE in 51H05 DN1p neurons diminished the width of summer E peak.

**Figure S4.**
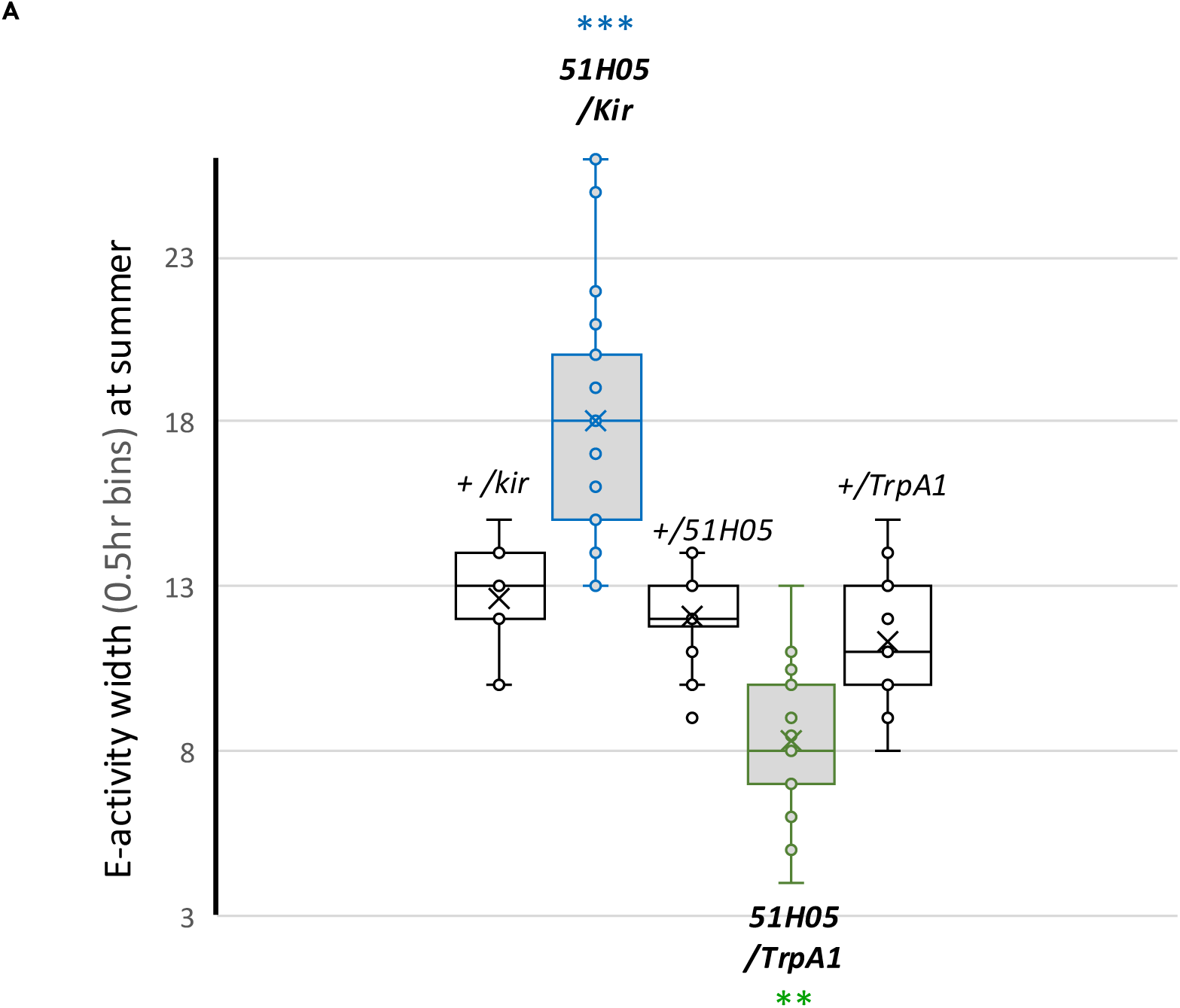
**A**. 51H05 neurons suppress locomotor activity. Targeted Kir2.1 expression in these neurons increased the width of E-activity in summer. On the other hand, TrpA1 expression in these neurons and elevating the temperature from 22°C to 30°C for 24 hours reduced the width of E-activity in summer.

**Figure S5.**
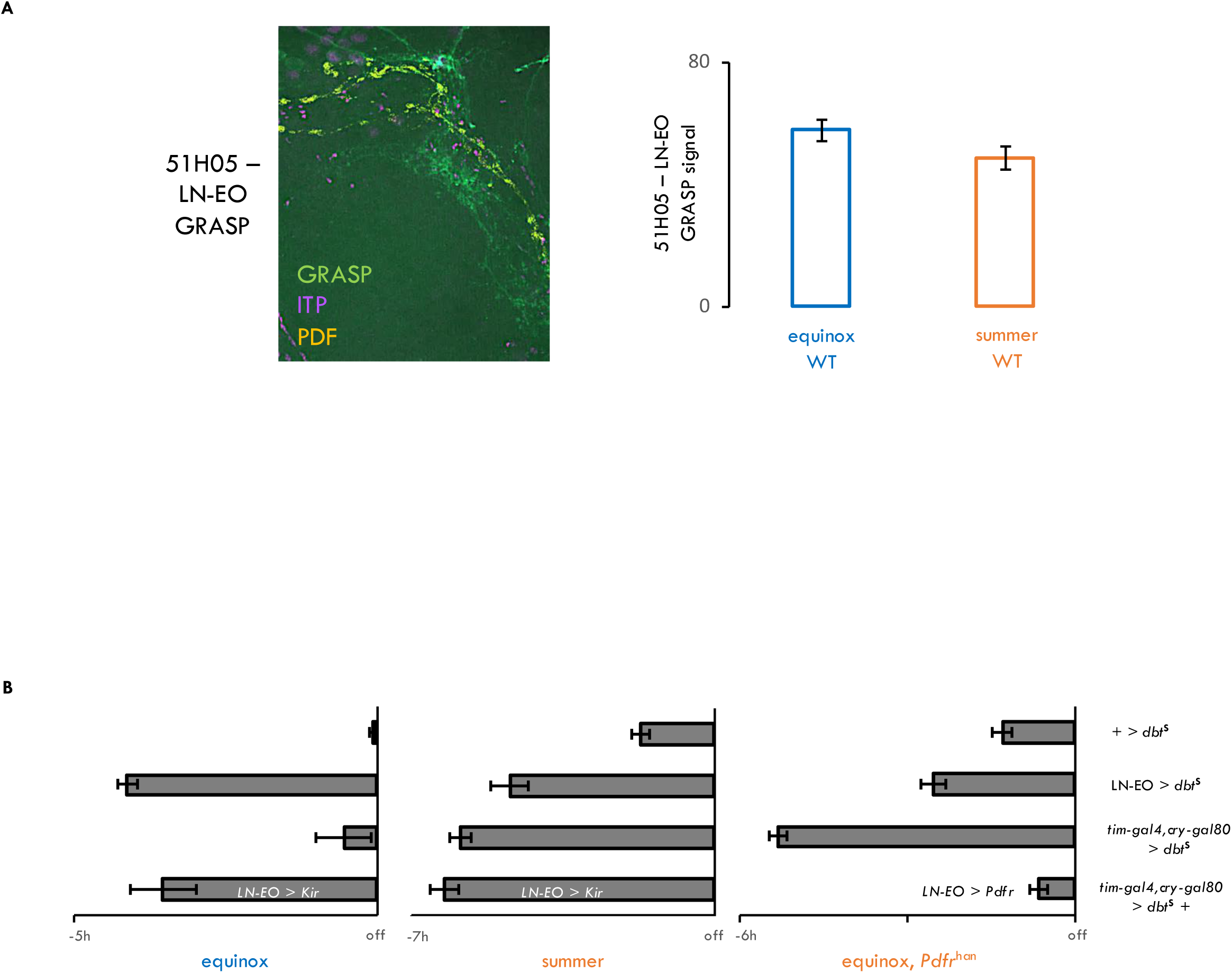
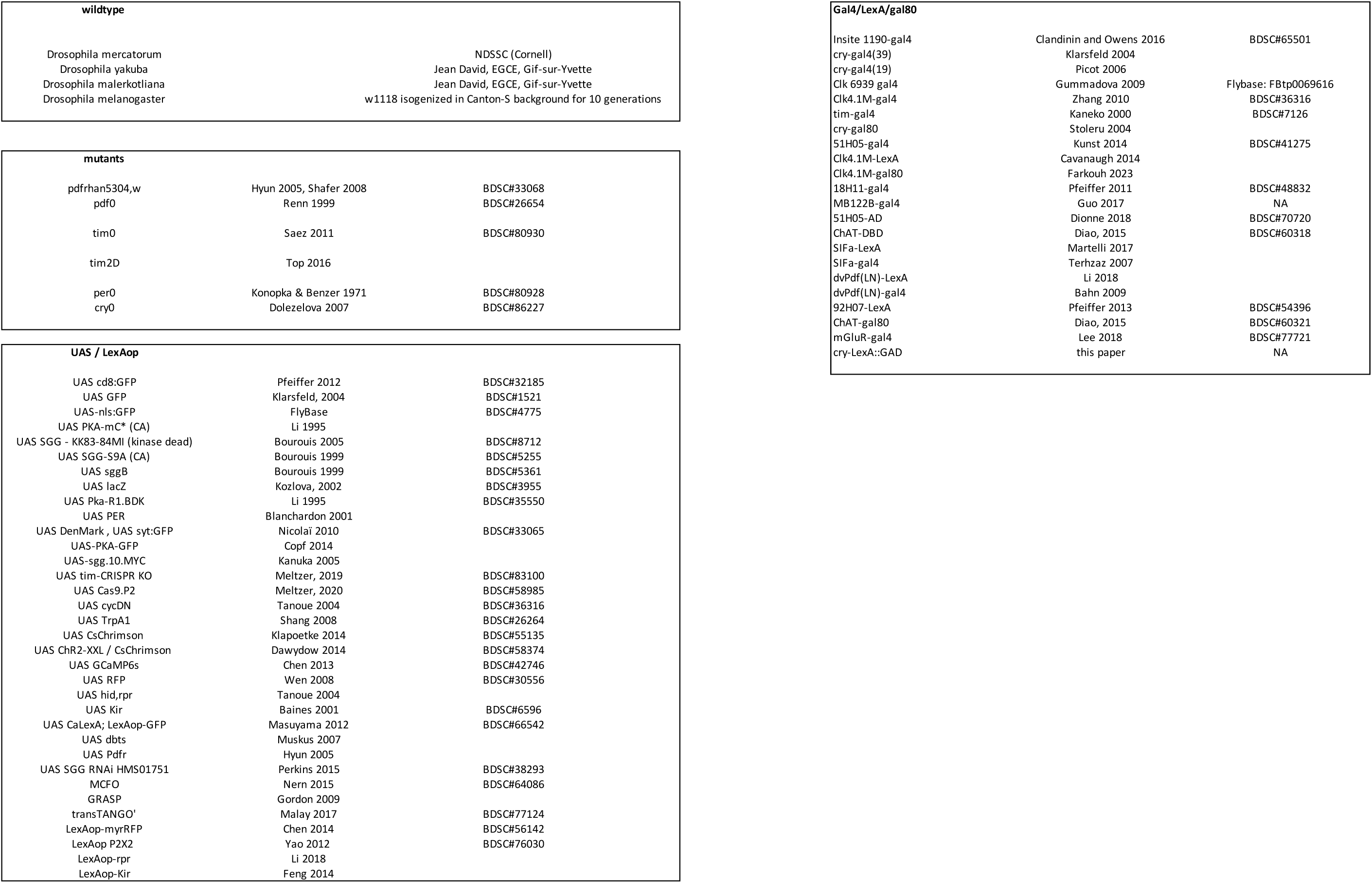
**A**. 51H05 DN1ps project ventrally toward the LNds. The strength of this anatomical connection does not vary according to the season. The GRASP signal between 51H05 DN1ps and LN-EO (ITP+ LNd and the 5^th^ sLNv) does not differ between equinox and summer. **B**. Seasonal structuring of the E-activity: The phase of the E-activity is regulated by the LN-EO in the ITP+ clock neurons. The 51H05 DN1ps achieve a conditional gain in their importance in phasing E activity in summer-like conditions or in absence of *Pdfr* in equinox or when the LN-EO was functionally removed from the circuit by expressing *Kir*. Additionally, *Pdfr* signaling in the LN-EO is crucial for designating context-dependent functional gain to the 51H05 DN1ps.

